# When Group Means Fail: Can One Size Fit All?

**DOI:** 10.1101/126490

**Authors:** Thomas H. Grandy, Ulman Lindenberger, Markus Werkle-Bergner

## Abstract

The present study examined whether a cognitive process model that is inferred based on group data holds, and is meaningful, at the level of the individual person. Investigation of this issue is tantamount to questioning that the same set and configuration of cognitive processes is present within all individuals, a usually untested assumption in standard group-based experiments. Search from memory as assessed with the Sternberg memory scanning paradigm is among the most widely studied phenomena in cognitive psychology. According to the original memory scanning model, search is serial and exhaustive. Here we critically examined the validity of this model across individuals and practice. 32 younger adults completed 1488 trials of the Sternberg task distributed over eight sessions. In the first session, group data followed the pattern predicted by the original model, replicating earlier findings. However, data from the first session were not sufficiently reliable for identifying whether each individual complied with the serial exhaustive search model. In sessions six to eight, when participants performed near asymptotic levels of performance, between-person differences were reliable, group data deviated substantially from the original memory search model, and the model fit only 13 of the 32 participants’ data. Our findings challenge the proposition that one general memory search process exists within a group of healthy younger adults, and questions the testability of this proposition at the individual level in single-session experiments. Implications for cognitive psychology and cognitive neuroscience are discussed with reference to earlier work emphasizing the explicit consideration of potentially existent individual differences.

## Introduction

“It is shortsighted to argue for one science to discover the general laws of mind or behavior and for a separate enterprise concerned with individual minds (…).” (Cronbach, 1957, p. 673)

This study examines whether a cognitive process model that is inferred based on group data holds, and is meaningful, at the level of the individual. Investigation of this issue is tantamount to questioning that the same set and configuration of cognitive processes is present within all individuals. Though this discussion has a long history (cf. Danziger, 1990), much of current research practice in experimental psychology and cognitive neuroscience tends to ignore its fundamental implications. Implicit to inferring cognitive processes from group data is— strictly speaking—the strong assumption and necessary precondition that the cognitive process under investigation is the same for every group member (cf. Caramazza, 1986; McCloskey & Caramazza, 1988). If this assumption does not hold, group data may represent a mixture of cognitive processes, and the cognitive process inferred may be in the worst case meaningless at the level of any individual (e.g., Hayes, 1953; Healey & Kahana, 2013; Miller, 2009; Molenaar & Campbell, 2009; Nesselroade, Gerstorf, Hardy, & Ram, 2007; Siegler, 1987).

As we will show in the present study, testing this assumption explicitly is possible but puts high demands on data reliability and stability. For the identification of a cognitive process at the individual level, the behavioral pattern found for a given individual has to be replicable on the person level, as unstable and varying processes are not easily separable from noise. The picture that emerges from the literature suggests that reliability of data from single measurement occasions is often insufficient to draw valid inferences about individual study participants (e.g., Carter, Krause, & Harbeson, 1986; Salthouse, Kausler, & Saults, 1986). From a technical point of view, sufficient reliability can be achieved by increasing the amount of data by pooling across multiple measurement occasions. However, from a psychological point of view, another question arises in this context: Can we assume the cognitive process under investigation to be stable across multiple measurement occasions, or does the cognitive process change—and if so, to what extent—when the same person is repeatedly confronted with a given task? This translates into two empirical questions, first, whether individuals would approximate stable, asymptotic performance after a limited amount of practice (across multiple measurement occasions), and second, whether the cognitive process under investigation changes as a function of practice (cf. Lövdén, Bäckman, Lindenberger, Schaefer, & Schmiedek, 2010).

On a more general level, the issues introduced above can be rephrased as a twofold inquiry about the generality of a cognitive process. On the one hand, we explore whether a cognitive process changes as a function of cumulative practice that is inherent to its repeated administration required to increase the amount of data. On the other hand, we examine whether a cognitive process that is assumed to be general is indeed discernible and equally found in all study participants.

In the following, we provide a short review of the history and issue of process identification at the group versus individual level, and discuss the role of reliability and stability as important prerequisites for making sound statements about individual people. We introduce and motivate the Sternberg paradigm (Sternberg, 1966, 1969, 1975) chosen as a target task for the general purpose of our inquiry, briefly summarize the discussion surrounding the original process model, and recapitulate previous findings regarding the reliability of model parameters and practice effects in the Sternberg task.

### Aggregate Level versus Individual Level

#### Historical Considerations

The research practice of inferring cognitive processes from aggregated data across individuals can be traced back to the beginnings of modern psychology. Here, we would like to direct attention to some lines of thought underlying the emergence of what nowadays appears as an unchallenged and undisputed research practice (see for a thorough treatment Danziger, 1987; Danziger, 1990; Gigerenzer, 1987a, 1987b; Lamiell, 2003).

Psychology has been often attested a fascination with classical physics (e.g., Gigerenzer, 1987a), arguably the leading discipline within the natural sciences. The emulation of classical physics as the prototypical natural science entailed the adoption of its scientific goal of finding general, nomothetic laws (Windelband, 1894, 1998/1894; see also Lamiell, 1998) for the workings of the mind, that is, laws that unequivocally and uniformly are common to all individuals (e.g., Hull, 1943; Krech, 1955). The assumption of the existence of a set of nomothetic laws for psychological processes was deeply incorporated into experimental psychological research; it defined what psychologists were searching for and at the same time, what they ignored. For instance, when considerable variance across individuals was observed in experimental settings, it was often attributed to a lack of control of boundary conditions in the experiment (cf. Gigerenzer, 1987a). An important achievement was the formulation of classical test theory, with its partitioning of observed variance into measurement error and true scores (Lord, Novick, & Birnbaum, 1968), a treatment of observational data that can be traced back to the beginnings of modern astronomy (cf. Stigler, 1986). However, by characterizing individual differences as ‘error’ around a single ‘true’ value, important hints towards individual differences were ignored and went unnoticed, contributing to the chasm of psychology into its “two disciplines” (Cronbach, 1957).

The advent, success, and appreciation of large-scale aggregate (social) statistics further promoted data averaging across individuals, as the attribute of social relevance and closeness to real life were important for the legitimization and promotion of the young scientific discipline of psychology. At the same time, the unit or level of investigation shifted and psychological claims were made by attributing psychological characteristics to collective rather than individual subjects (Danziger, 1990). The distinction between the collective and the individual has been blurred from the very beginning, as reflected, for instance, in the indifference of Thurstone against interchanging intraindividual (1927b) with interindividual (1927a) variability (see Danziger, 1990; Gigerenzer, 1987a, 1987b). Not differentiating between the fundamental implications of ascribing the ‘error’ to intraindividual versus interindividual variance constituted a highly critical step in the shift from individuals to aggregates without adaptation of the scope of validity of the statements and laws derived (cf. Kraemer, 1978; Molenaar, 2004; Voelkle, Brose, Schmiedek, & Lindenberger, 2014).

Taken together, the history of experimental psychology has been marked by a search for general, nomothetic laws. In conjunction with the success of aggregate statistics (e.g., analysis of variance), psychology as a science of the laws that govern individual minds has been largely left behind (Danziger, 1987, 1990; Gigerenzer, 1987a, 1987b; Lamiell, 2003; Molenaar, 2004; Nesselroade, 2010).

#### Criticism of the Practice of Aggregating

The historical spotlights presented above are neither exhaustive nor sufficient to explain what has been coined “the triumph of the aggregate” (Danziger, 1990); rather, they are influential lines of thought that have played their role in the development of current research practices in mainstream experimental psychology and cognitive neuroscience. At the same time, the practice of aggregating data across individuals has been continuously challenged and criticized.

Hayes’ (1953) analysis of learning trajectories provided a prominent case for questioning the practice of replacing individual by aggregated data. When performance improves in the form of a step function, and the steps occur at different times during learning for different individuals, aggregated data would suggest a sigmoid-like learning trajectory instead of a step function (see also Lewandowsky & Farrell, 2011). Thus, the two levels lead to distinctly different interpretations of the learning process. In a similar vein, aggregation of individual exponential curves may lead to a power curve in the aggregate (e.g., Anderson & Tweney, 1997; Estes, 1956; Heathcote, Brown, & Mewhort, 2000).

Beyond the learning literature, the importance of individual differences has, for example, been emphasized with regards to differential strategy use in memory or arithmetic tasks (e.g., Bailey, Dunlosky, & Hertzog, 2009; Dunlosky & Hertzog, 1998; Healey & Kahana, 2013; Miller, Donovan, Bennett, Aminoff, & Mayer, 2012; Siegler, 1987), and it has been pointed out, that uncritical aggregation of data at worst results in a meaningless mixture of strategies and corresponding cognitive processes. In addition, not only in complex cognitive tasks substantial individual differences have been reported. Individual differences have also been described as a challenge for lawful psychophysical relationships derived from aggregated data (e.g., Gigerenzer, 1987a; Gigerenzer & Strube, 1983; Reynolds & Stevens, 1960).

On the other hand, cognitive neuropsychology traditionally has dealt with the investigation of single patients. By means of single-case studies, it has provided some major contributions to the understanding of the relationship between cognitive functions and brain structures, for instance, in uncovering the fundamental role of the hippocampus in episodic memory (Scoville & Milner, 1957). Nonetheless, there also has been an extensive debate on the significance and weight of single-case studies as compared to group studies (cf. Caramazza, 1986; Caramazza & McCloskey, 1988; McCloskey, 1993; McCloskey & Caramazza, 1988; Robertson, Knight, Rafal, & Shimamura, 1993). McCloskey and Caramazza (1988) stressed the appreciation of individual performance patterns in the investigation and understanding of pathological conditions of the brain. In contrast, Robertson and colleagues (1993) emphasized the difficulties in integrating evidence from isolated cases, and promoted group studies as being more apt for the examination of functionally distinct components or modules of cognition. However, the controversy seems to have been mainly circling around the proper treatment of patient data, whereas homogeneity of cognitive processes in normal individuals is commonly a priori taken for granted by assuming that the functional architecture of the cognitive system is the same for all normal individuals (“assumption of universality”; Caramazza, 1986; McCloskey & Caramazza, 1988). Only recently, it has been pointed out that “relatively little work has been done on understanding the extent and nature of individual variability with regard to the types of cognitive mechanisms commonly investigated in cognitive psychology and neuropsychology” (Rapp, 2012, p. 8).

Finally, the issue of inferring psychological structures from analyzing group versus individual data has also (re-)gained attention in differential and developmental psychology (cf. Hamaker, Dolan, & Molenaar, 2005; Lamiell, 2003; Molenaar & Campbell, 2009; Molenaar & Newell, 2010; Nesselroade, 2001, 2002, 2004, 2010; Nesselroade et al., 2007; Rogers, Hertzog, & Fisk, 2000; Voelkle et al., 2014), with some reference to ergodicity theory (Molenaar, 2004; Molenaar & Campbell, 2009). Here, some of the arguments put forward are reminiscent of the description of the “ecological fallacy” (Robinson, 1950), which implies that incorrect conclusions about the interrelation of characteristics of individuals are derived when calculating the correlation on aggregated data. Of note, Kraemer (1978) has convincingly elaborated that not aggregation or the level of aggregation per se may be critical, but that great care has to be taken that the interpretation of findings does not refer to a level that was not subject to investigation or analysis.

#### Prerequisites for Inquiries at the Individual Level

Assessing psychological processes at the individual level requires a sufficient level of reliability and stability at the within-person level. Increasing data density through repeated measurements is an effective way of enhancing reliability. Implicitly, this can be observed in the early days of experimental psychology, where the unit of investigation was the individual and researchers often served as their own subjects (Danziger, 1987, 1990). As a case in point, Ebbinghaus (1885) went through an impressive amount of extensive repetitions of learning arbitrary verbal material to derive his principles of learning and forgetting. The practice of extensive repeated assessments of individuals has continued up to today in some areas of psychology, such as mathematical psychology and formal modeling, where high data density is required (e.g., Ashby, Tein, & Balakrishnan, 1993; Donkin & Nosofsky, 2012; Nosofsky, Little, Donkin, & Fific, 2011; Ratcliff, 1978; Roberts, 2004; Townsend & Fifić, 2004), and generality of a process is often inferred from replicability of a model across several individuals. However, sample sizes in these studies are often small, thereby not allowing for a systematic investigation of the extent and nature of individual differences.

From the perspective of classical test theory (Lord et al., 1968), the issue of reliability and stability can be recast as a problem of the relationship between the signal of interest (true value) and noise (error). Increasing data density leads to an increase of the signal-to-noise ratio and hence increased reliability (see also W. Brown, 1910; Spearman, 1910). Ideally, the signal-to-noise ratio increases proportional to the square root of the number of measurements. However, preconditions are that (1) signal and noise are uncorrelated, (2) the signal is constant in the replicate measurements, and (3) the noise is random, with a mean of zero and constant variance in the replicate measurements (cf. van Drongelen, 2006). However, these preconditions are rarely met in repeated assessments in psychology. Due to effects of, for example, learning, practice, or memory, repeated measurements of individuals often violate the assumption of independence of errors (see also Gigerenzer, 1987a). Furthermore, the independence requirement is less likely to be met when studying psychological phenomena in naïve research subjects, as is typically done today (cf. Danziger, 1990). In the case of early self-experimentation, researchers were often highly acquainted with their tasks and proficient in accomplishing the tasks they investigated; thus, learning, practice, or memory effects were more likely negligible.

When investigating more than a handful of individuals, one can rarely draw upon individuals experienced and skilled in specific experimental paradigms. Thus, we were interested in creating an experimental situation were a larger sample of younger adults would approach stable, asymptotic performance within a limited number of sessions. Here, we assessed stability explicitly by testing (1) whether performance reached asymptotic levels within limited time (mean stability), and (2) whether interindividual differences across sessions stabilized (relative stability; cf. Kagan, 1980). High mean stability together with relative stability is the necessary precondition for enhancing the reliability of measurements by means of data aggregation at the individual level. In order to approximate the amount of ‘true’ noise versus systematic fluctuations, for instance, due to practice, we formally tested whether reliability increased as a function of increasing the number of measurement occasions as predicted by the Spearman-Brown formula. With respect to validity, we assessed whether the same (or at least similar) cognitive processes were at work when completing the task after practice as compared to the first encounter with the task.

### The Sternberg Paradigm

The above questions were investigated with the Sternberg memory scanning paradigm (Sternberg, 1966, 1969, 1975). We have chosen this paradigm in order to make solid contact to earlier research. The Sternberg paradigm is well established, and the range of phenomena that may be observed within this paradigm have been extensively investigated and discussed. Even though the Sternberg paradigm is one of the classic paradigms in experimental psychology, debates about the nature of relevant search processes remain to be resolved (e.g., Ashby et al., 1993; Donkin & Nosofsky, 2012; Nosofsky et al., 2011; Ratcliff, 1978; Sternberg, 1975; Townsend & Fifić, 2004). The paradigm is also frequently adopted in cognitive neuroscience (e.g., Ole Jensen, Gelfand, Kounios, & Lisman, 2002; O. Jensen & Tesche, 2002; Näpflin, Wildi, & Sarnthein, 2008; Obleser, Wostmann, Hellbernd, Wilsch, & Maess, 2012; Pelosi, Hayward, & Blumhardt, 1995, 1998; Pratt, Michalewski, Barrett, & Starr, 1989; Pratt, Michalewski, Patterson, & Starr, 1989a, 1989b; Tuladhar et al., 2007). Thus, the Sternberg paradigm continues to be a tool for research on the organization of search processes in memory. Our contribution to this research consists in questioning the validity of only one memory search model for the Sternberg paradigm (“assumption of universality”; Caramazza, 1986) by re-introducing the individual level of observation and analysis. To the best of our knowledge, this has not yet been done with the necessary number of repeated observations within individuals and at the same time with a larger number of individuals.

In the standard version of the Sternberg paradigm, participants are presented sequentially or concurrently with a set of *n* distinct items (e.g., digits) that are to be held in memory. Set size *n* varies between one and six items, and the items that are held in memory are called the *positive set*; items of the overall item pool not contained in the positive set constitute the *negative set*. The overall item pool is limited in number, for example, comprising the digits zero to nine. After a short interval of one or two seconds, a probe item is presented, that either belongs to the positive set (*positive probe*), or to the negative set (*negative probe*). Participants are instructed to indicate as fast and accurately as possible whether or not the probe was a member of the memorized set by pressing one of two alternative buttons or levers (*positive* or *yes response*, if the probe was part of the positive set, and *negative* or *no response* if it was not).

Two variants of the Sternberg paradigm exist regarding the to-be-memorized set of items. In the *varied-set procedure*, a new set of *n* items is presented in every trial, followed by a single probe item. In the *fixed-set procedure*, presentation of one set of items is followed by several probes, for example 120 successive probe trials for the same positive set (cf. Sternberg, 1966). In the current study, we employed the varied-set procedure.

The most fundamental findings of the original set of experiments conducted by Sternberg (cf. 1966, 1969, 1975) were (a) a linear increase of mean response times (RTs) to the probes as a function of set size *n*, and (b) the equality of the slopes for the *yes* and *no* RT functions. Sternberg concluded that the findings strongly supported a *serial* and *exhaustive memory search* process. Serial memory search was inferred from the linearity of slopes; for every additional item in memory, a fixed amount of additional time for the memory search was required.

Exhaustiveness of memory search was deduced from the equality of slopes; equal slopes indicate that the cumulative search time for positive and negative probes is the same, thus, the complete memorized set is scanned before the response is initialized.

The generality of these original findings has been questioned for long. In the following, we introduce some of the divergent findings and claims in relation to the question whether there are distinct individual differences in memory search process. We then briefly review the literature regarding reliability of slopes as well as practice effects in the Sternberg paradigm.

#### Controversies about the Memory Search Process in the Sternberg Paradigm

The main findings of the Sternberg paradigm, namely linear and equal increase of RTs as a function of set size and probe type, and their interpretation as serial exhaustive memory search were replicated several times (e.g., Burrows & Okada, 1973; Chase & Calfee, 1969; Harris & Fleer, 1974; McCauley, Kellas, Dugas, & DeVellis, 1976; Swanson, Johnsen, & Briggs, 1972; Wingfield, 1973; Wingfield & Branca, 1970). Nonetheless, there has been a long-standing discussion regarding the generality of the original findings that have put the serial exhaustive memory search model into question. In this paper, we focus on *serial self-terminating memory search* as the most prominent alternative to exhaustive memory search (cf. Sternberg, 1969, 1975). Here, in contrast to serial exhaustive memory search, the search process stops once a positive probe has been found in memory. As discussed by Sternberg, the slope ratio between the slopes of *yes* and *no* responses allows differentiation of serial exhaustive and self-terminating memory search. As described above, a slope ratio of 1.0 (equal slopes) can be seen as indicating serial exhaustive memory search. On the other hand, a slope ratio of 0.5 may indicate self-terminating memory search, as a match for positive probes should be found on average, after half the memorized list has been searched. Some studies reported subgroups or individuals showing evidence for a self-terminating memory search process (Clifton & Birenbaum, 1970; Corballis, Kirby, & Miller, 1972; Corballis & Miller, 1973; Klatzky & Atkinson, 1970; Swinney & Taylor, 1971; see also Sternberg, 1975). Furthermore, at a closer look, in some studies, slope ratios—re-calculated from reported slopes—were found to be below 1.0 (e.g., Chase & Calfee, 1969; Swanson et al., 1972), raising the question whether memory search processes differed among individual participants. In an interesting recent study, cognitive strategies reported by individuals varied widely (Corbin & Marquer, 2009); however, no relationship between reported strategies and RT patterns was found. From a measurement perspective, the imperfect reliability of slopes reported in the literature (see below) raises the question whether slope ratios collected within a single session can be unambiguously ascribed to individual participants. In the study by Klatzky and Atkinson (1970), for example, individual slope ratios were broadly distributed from 0.03 to 2.63.

#### Reliability and Stability of the Memory Scanning Rate

Low reliability and stability of slopes has been discussed as a major impediment for the use of the Sternberg paradigm in the assessment of individual differences (Carter, Kennedy, Bittner, & Krause, 1980; Carter et al., 1986; Roznowski & Smith, 1993). Furthermore, low reliability of slopes makes the direct assessment of individual differences in the slope ratio—as an indicator of competing memory search processes like exhaustive versus self-terminating—difficult, as the contribution of slope unreliability is amplified in the calculation of the slope ratio.

Split-half reliabilities of slopes have been reported to range between .69 and .83 (H. L. Brown & Kirsner, 1980; Chiang & Atkinson, 1976). Within-session stability across blocks of trials were found to range between .47 and .52 (Longstreth & Madigan, 1982), whereas values for stability across sessions have been reported to range from −.57 (!) to .78 (Carter et al., 1980; Carter et al., 1986; Chiang & Atkinson, 1976; Roznowski & Smith, 1993). However, for all studies thus far, only a low number of trials per condition (set size × probe type) was assessed, ranging from approximately three to eight trials per condition in the stability studies (Carter et al., 1980; Longstreth & Madigan, 1982; Roznowski & Smith, 1993), and consisting of approximately 30 trials per condition in the reliability studies (H. L. Brown & Kirsner, 1980; Longstreth & Madigan, 1982). To the best of our knowledge, the reliability of slope ratios has not been reported up to now, despite reports on individual differences in the slope ratio (Clifton & Birenbaum, 1970; Corballis et al., 1972; Corballis & Miller, 1973; Klatzky & Atkinson, 1970; Swinney & Taylor, 1971).

#### Practice in the Sternberg Paradigm

Several studies have investigated the effect of extended practice on performance with the Sternberg paradigm. The large majority used the fixed-set procedure, that is, the same digits comprised the positive set within one (Kristofferson, 1972a) or even throughout all sessions (Kristofferson, 1972b, 1977; Lively, 1972; Ross, 1970; Simpson, 1972). Thus, with the exception of Kristofferson (1972a), responses in the above cited studies were consistently associated with specific stimuli. Within all studies, a reliable change of the zero intercept of the RTs as a function of set size was observed. In contrast, evidence for a decrease of the memory scanning rate was mixed; apart from the study by Kristofferson (1972a) where responses were not associated with specific stimuli, most studies reported reliably decreasing slopes as a function of practice.

We are aware of only two studies (Carter et al., 1980; Nickerson, 1966) that tested practice effects with the varied-set procedure of the Sternberg paradigm. In the study by Nickerson (Nickerson, 1966), intercepts were found to decrease but slopes did not change with practice. However, the task used by Nickerson (1966) deviated in various aspects from the original Sternberg paradigm: only set sizes of one and four were employed, precluding the assessment of linearity of slopes, stimuli were presented concurrently, the retention interval only lasted one second, and letters instead of digits were used as stimuli. In contrast, Carter and colleagues (1980) found no change of the intercept with practice, but a substantial decrease in the slope. Thus, comprehensive and consistent evidence on the stability of the RT effects for the varied-set procedure of the Sternberg paradigm is lacking in the context of extensive practice.

As a note aside, in the two studies that reported memory scanning rates not to be affected by practice (Kristofferson, 1972a; Nickerson, 1966), the scanning rates were reported to range between 33 and 36 ms/item. The fact that these scanning rates did not differ largely from the original findings by Sternberg (1966; 38 ms/digit), and that slopes were found to be equal for *yes* and *no* responses and did not change by practice were taken as evidence supporting the strong notion of a general memory scanning rate for digits (cf. Cavanagh, 1972; Sternberg, 1975).

### Goals of the Present Study

Using the classic Sternberg paradigm (Sternberg, 1966, 1969, 1975), the main goal of this study was to test in a specific instance the standard and often implicit assumption that cognitive processes are general and identical within and across healthy individuals (cf. Caramazza, 1986; McCloskey & Caramazza, 1988). We tested this assumption in a sample of 32 healthy adults aged 20 to 27 years with similar levels of education. As introduced, the study’s main goals entailed a set of interrelated issues that are listed below.

#### Mean Level: Replication of the Sternberg Effect

We started by investigating the replicability of the main findings of the original varied-set Sternberg paradigm (1966) at the aggregate level. This was done primarily to assure that the task implementation was not deviating substantially from the original paradigm, and to provide a sound starting point for subsequent investigations.

#### Mean Level: Practice and the Sternberg Effect

We investigated practice effects for several reasons. Primarily, it was of interest to find out whether a sample of young adults would approach asymptotic performance within a limited amount of time. As mentioned before, mean level stability of data—together with relative stability—is an important precondition for meaningful aggregation of data across sessions with the goal of achieving sufficient data density at the individual level to increase reliability. We operationally defined performance close to the asymptote when changes in mean RTs from session to session would no longer differ reliably from zero at the group level, resulting in stable performance at the aggregate level. In addition, we aimed at testing whether the memory search model would apply equally well before and after practice, that is, generalized to the performance after practice inherent in the repeated assessments of the task. Specifically, we were interested in examining whether and to what extent slopes for *yes* and *no* responses as well as their ratio, characterizing the memory search process, would remain invariant or change with practice.

#### Reliability and Stability of Data

As introduced above, the key to investigating psychological processes at the individual level is high reliability of measurements (and derived parameters). With respect to the Sternberg paradigm, the unreliability of slopes has imposed a fundamental limitation on the detailed investigation of individual differences in memory search processes, as the effects of unreliability accumulate when forming ratio scores (cf. Carter et al., 1980; Carter et al., 1986). Previous studies have only investigated individual differences in slopes and their reliability with restricted numbers of trials. Here, we attempted to assess the dependency of reliability on the number of trials acquired across consecutive sessions.

High relative together with mean level stability across sessions imply a minimization of systematic changes in the repeated measurements that occur, for example, because of differential practice effects, and constitute the necessary preconditions for data aggregation with the goal of systematically increasing reliability. Hence, we examined whether the structure of individual differences changed with practice, and whether asymptotic performance coincided with a high level of relative stability. Within session split-half reliabilities provided an estimate for the upper bound of relative stability. Specifically, we tested whether reliability increased as predicted by the Spearman-Brown formula when pooling data across sessions, thus, denoting optimal ‘noise’ reduction and validating the approach to acquire data from multiple sessions in a principled way in order to obtain a sufficient reliability for the identification of memory search processes at the individual level.

Of note, from the perspective of testing cognitive process models at the individual level, measuring reliability and stability (across individuals) can be seen as an auxiliary strategy for extrapolating the replicability of parameters at the individual level. The adequacy of this rationale relies on the presence of substantial between-person differences which indeed have been suggested by the above reviewed literature (see also Hunt, Frost, & Lunneborg, 1973; Keating & Bobbitt, 1978).

#### Individual Level

The main goal of this study was to capture individual memory search behavior under conditions of enhanced reliability. We aimed to investigate whether the behavior of individuals within a group of healthy younger adults is well described by a model of serial and exhaustive memory search once they have reached levels of performance that provide sufficient data for identifying memory search parameters at the individual level.

In concert, our analyses were targeted at scrutinizing the generality and validity of a prominent process model in cognitive psychology. If model parameters support the assumption of serial exhaustive search for some individuals but not for others, or if slope ratios change with practice, then one would need to conclude that group-based analyses of single-session Sternberg data yield a picture of process generality that may not correspond to the processing realities of individual people and their performance when being acquainted with the cognitive task at hand.

## Methods

### Participants

The sample comprised 32 young adults (*M_age_* = 23.3 years, *SD* = 2.0, range 19.6 to 26.8 years; 17 women), the majority being university students (*n* = 28). The participants were recruited from the participant pool of the Max Planck Institute for Human Development, Berlin, Germany (MPIB). All participants gave written informed consent according to institutional guidelines of the ethics committee of the MPIB. Participants were right-handed, as assessed with a modified version of the Edinburgh Handedness Inventory (Oldfield, 1971), and had normal or corrected-to-normal vision, as assessed with the *Freiburg Visual Acuity test* (Bach, 1996, 2007). Participants reported to be in good health with no known history of neurological or psychiatric disease, and were paid for participation (8.08 € per hour, 25.00 € for completing the study within 16 days, and a performance-dependent bonus of 28.00 €, see below).

### Study Design

The study consisted of one *covariate session*, followed by eight *repeated sessions* in which the Sternberg task and three additional cognitive tasks were administered (Figure 1; Appendix A); in the following, we refer to the eight repeated sessions as *sessions one to eight*. All sessions were completed on consecutive days with exception of Sundays and session eight. The covariate session and session one, as well as sessions six and seven were always completed on immediately following days. Session seven was completed seven days after session one by all but one participant (eight days, due to one break of two days). Session eight was conducted approximately one week after session seven (*M* = 7.3 days, *SD* = 1.4), and served to estimate the stability of practice gains across several days. In sessions one, seven, and eight electroencephalogram (EEG) was recorded concurrently with the Sternberg and a Choice Reaction task.

**Figure 1.**
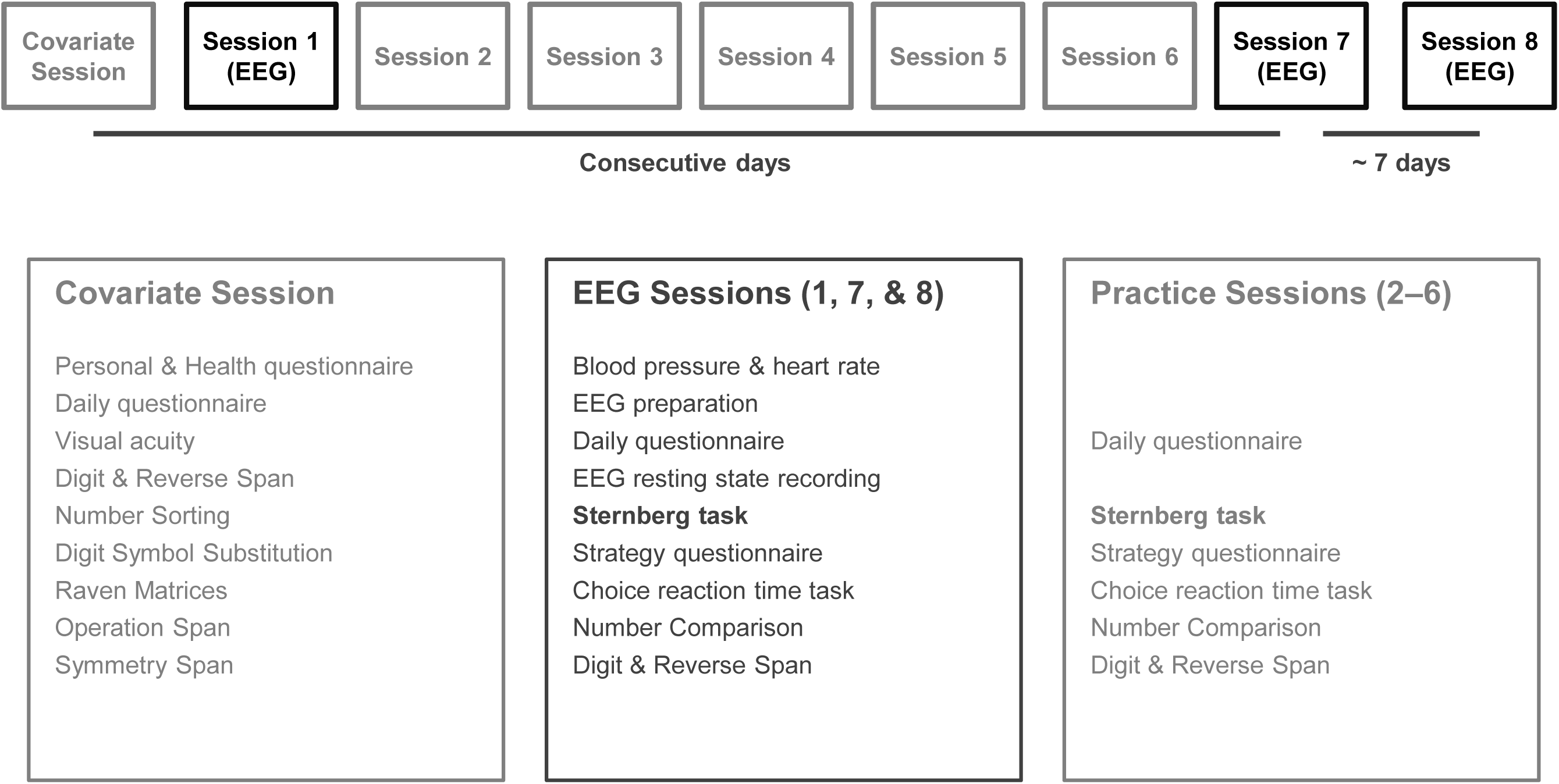
Overview of study design and listing of tasks and questionnaires employed. A short description of tasks is provided in Appendix A.

In the covariate session (2.5–3.0 h) questionnaires and several tests with a focus on short-term and working memory tasks were completed. In sessions one to eight four tasks, including the Sternberg task, were assessed repeatedly. The sessions with concurrent EEG lasted approximately three and a half to four hours, including approximately one and a half hour of EEG preparation. The sessions without EEG (i.e., sessions two to six) lasted approximately one and a half to two hours. Given that this article focuses on the Sternberg paradigm, we will only describe the Sternberg task in detail here; a brief description of all tasks used in this study is provided in Appendix A.

### Sternberg Task

The version of the Sternberg task used here closely followed the original varied-set procedure by Sternberg (1966). Digits zero to nine were used as stimuli. To maximize the number of trials per set size and probe type, we only assessed set sizes of two, four, and six. The participants initiated the task by pressing first the right and then the left response key with their respective index fingers; this way we ensured that participants had their index fingers on the correct response keys upon start of each trial. Digits were presented consecutively every 1.2 sec. They were displayed on the screen for 0.2 sec, followed by a blank inter-stimulus interval (ISI) of 1.0 sec. After presentation (offset) of the last to be memorized digit, a 3.0 sec delay (retention interval) preceded the presentation of the probe digit, corresponding with the 2.0 sec delay in Sternberg (1966), where digits were presented for 1.2 sec without ISI. Presentation of the probe-item also lasted for 0.2 sec, followed by a blank screen for 2.0 sec in which the response of the participants was recorded. A beep tone indicated the end of the trial, and a message on the monitor prompted the participants to verbally recall the to-be-memorized set of digits in the correct serial order of their presentation. Serial recall of all trials was recorded digitally.

Half of the probes were *positive* probes (i.e., were contained in the to-be-memorized set), and half of the probes were *negative* probes (i.e., were not contained in the to-be-memorized set). To be memorized digits were randomly drawn without replacement from the ten digits. The ten digits were equally distributed across positive as well as negative probes, and across serial positions in the to-be-memorized sets. Each serial position was probed equally often. For each combination of set size (2, 4, 6) × probe type (positive, negative), 31 trials were carried out, leading altogether to 186 trials per session. The six set size × probe type combinations were randomly distributed across four blocks (the first block consisting of 48 and the second to fourth block of 46 trials); every participant obtained a different randomization of the order of stimuli, set sizes, and probe types. Participants were not told in advance which set size the following trial would comprise. In session one, participants additionally practiced the Sternberg task with 24 trials (four trials per set size and probe type).

Each block ended with a summary feedback of the overall mean RT and accuracy within the current session up to that point. To encourage high motivation and performance throughout the whole study, a bonus (28.00 €) was paid under the following conditions: overall mean RT had to be faster or at least equal to the overall mean RT of the preceding session, while overall accuracy had to remain higher than 90%.

Participants were seated at a distance of 80 cm (eyes to center of the monitor) in front of a LCD monitor with a refresh rate of 60 Hz. Digits extended approximately 2.5° of visual angle in the vertical and 1.8° of visual angle in the horizontal direction, and were presented in white on a black background. Stimulus-presentation and recording of behavioral responses were controlled with E-Prime 2.0 (Psychology Software Tools, Inc., Pittsburgh, PA, USA).

### Statistical Analyses

Statistical analyses were conducted using MATLAB (The MathWorks Inc., Natick, MA, USA), SPSS (SPSS Inc., Chicago, IL, USA), R (R Core Team, 2013; http://www.R-project.org/), and SAS PROC MIXED (SAS Institute Inc., Cary, NC, USA). Only response times (RTs) of correct responses were retained for analysis. Within each of the six conditions (set sizes × probe types), correct RTs more than ± 2.5 *SD*s from the individual mean RTs were excluded recursively from the analysis, leading to on average of 1.6 discarded correct responses per condition.

#### Model Comparisons

Throughout this report we fitted mean RTs across three set sizes and two probe types with an unconstrained linear model, allowing for slopes and intercepts to be different across the two probe types (*positive* and *negative*, or *yes* and *no* responses). This unconstrained model was compared to a constrained model where slopes were restricted to be equal. The latter model effectively represents the exhaustive memory scanning model introduced by Sternberg (1966, 1969, 1975). Using two specified models allowed us to statistically test for equality of slopes, as described below.

Following common practice (cf. Sternberg, 1966), we first fitted linear regression models to the overall mean RTs. However, this practice only provides limited statistical power for formal model comparisons, as models were fitted to only six overall mean RTs (set size × probe type). Thus, model comparisons were also conducted within the framework of multilevel (mixed-effects) modeling (MLM) using SAS PROC MIXED (cf. Singer, 1998; Snijders & Bosker, 1999). Fixed effects were modeled equivalently to the two models introduced above. Instead of fitting overall mean RTs only, RTs were now allowed to vary across individuals (random effects). Parameters were fitted using maximum likelihood (ML) estimation with an unconstrained variance-covariance matrix, allowing all variances and covariances to be estimated freely. With this MLM specification, the fixed effects are identical to the regression parameters of the fit to the overall mean RTs in the unconstrained model. Within MLM, we tested equivalence of parameters across models by means of the likelihood ratio test. Hereby, equivalence of parameters, for example slopes of *yes* and *no* responses, is tested by constraining these parameters to be equal and by testing whether the resulting change in −2 log likelihood (denoted as Δ*χ*^2^) is reliable, with the degrees of freedom equal to the number of constrained parameters (cf. Bollen, 1989; Kline, 2005).

#### Practice Effects

Overall practice effects were assessed by means of repeated measures analysis of variance (rmANOVA), with factors session (1–8) × set size (2, 4, 6) × probe type (positive, negative). Whenever the assumption of sphericity was violated (Mauchly’s test), Greenhouse-Geisser corrected degrees of freedom and *p* values are reported. Session by session comparisons of mean RTs within set sizes and probe types were conducted using paired-sample Student’s *t*-tests. Asymptotic performance at the mean level was operationally defined as statistically non-reliable differences between sessions. Note that not correcting for multiple comparisons lead to a more conservative estimate of asymptotic performance. Mean stability in sessions six to eight was in addition confirmed by an omnibus post-hoc rmANOVA with factors session (6–8) × set size × probe type.

#### Reliability and Stability Analyses

Reliability and stability analyses served to identify conditions under which memory search processes can be studied at the individual level. We first assessed split-half reliabilities of mean RTs, intercepts, slopes, and slope ratios within sessions one to eight. Since RTs show considerably broad distributions *within* individuals relative to between-person differences in mean RTs—in contrast to the psychometric assumption of homogeneous response tendencies at the individual level—, any split of the RTs into two halves within individuals may provide relatively arbitrary reliability estimates. Thus, in order to get stable estimates, we repeated calculation of split-half reliabilities with 1,000 random splits of RTs within individuals and conditions, and report the mean split-half reliability of 1,000 random splits. All reported averages of correlation coefficients were obtained by applying the Fisher *Z*-transformation to correlation values prior to averaging, and subsequently applying the inverse transformation to the averaged *Z* value (cf. Silver & Dunlap, 1987). To provide a general picture of reliability, and given that the main focus of this article is on the slope ratio, average reliabilities of intercepts and slopes of *yes* and *no* responses are reported here.

Next, we assessed the relative (i.e., rank order) stability across sessions. Here, correlation coefficients between mean RTs, intercepts, slopes and the slope ratios of all pairs of sessions are reported. Stability coefficients reported for intercepts and slopes are again averaged across *yes* and *no* responses. In order to compare the average stability during initial practice to the stability after some practice, stability coefficients between sessions one and four were compared to stability coefficients between sessions five and eight with Student’s *t*-tests. Furthermore, mean stability of sessions five to eight was also compared to the mean stability between distant sessions, that is, between sessions with at least three sessions in between, with Student’s *t*-tests. *t*-tests were performed on Fisher *Z*-transformed correlation coefficients. *F*-tests indicated equal variances for all comparisons [*F*s(5,5) ≤ 5.17, *p*s > .096; *F*s(9,5) ≤ 2.29, *p*s > .376]; Kolmogorov-Smirnov tests did not indicate reliable deviations from normal distribution for all *Z*-transformed correlation coefficients tested (*Z*s ≤ 0.27, *p*s > .712).

Finally, we tested whether increasing the amount of data by pooling data across sessions would lead to predictable increases in split-half reliability, denoting optimal ‘noise’ reduction. Reliability coefficients for pooled sessions were obtained as described above, with RTs from two and three neighboring sessions being pooled prior to randomly splitting the RTs into two halves. We focused on the comparison of sessions one to three with sessions six to eight. Reliability values reported are the average reliability of single ore pooled sessions, respectively. Predicted reliabilities as a function of the amount of pooled data were estimated based on the average single session reliabilities using the Spearman-Brown prediction formula. Comparisons of observed and predicted correlation coefficients are provided as the deviation of the observed from the predicted correlation coefficients expressed as *z*-values.

Throughout this report, we provide Spearman rank correlation coefficients (*ρ*), in order to obtain a robust estimate of reliability and stability, not being as sensitive to outliers. Equivalent analyses not reported here using Pearson correlation coefficient showed the same pattern of results.

#### Individual Level Analyses

Analyses at the individual level were performed by fitting the unconstrained model to mean RTs at asymptotic performance (sessions six to eight) within every participant. An estimate of the stability of the slope ratio as the main indicator for process identification was obtained by bootstrapping the mean RTs within each condition and subsequent least-squares fit with the unconstrained model (10,000 re-samplings per participant). Bootstrapped 95% confidence intervals (CIs) for the slope ratios of all 32 participants are reported. In order to compare the memory search process identification at asymptotic performance with conventional single session data, model fitting and bootstrapping of the slope ratio was conducted in the same way with data from session one.

## Results

### Mean Level Perspective

#### Replication of Sternberg’s Original Findings in the First Session

In the first session, the core findings of the Sternberg paradigm were replicated: (1) linear increase of RTs as a function set size (items in memory) and (2) equality of slopes of *yes* and *no* responses (Figure 2). Accuracies were found to be on a sufficiently high level (i.e., above 90 %; cf. Sternberg, 1975). Participants committed on average approximately one error per set size and response type (accuracy 96.6 to 98.3%) with the exception of *yes* responses in set size six (93.4%, i.e., on average approximately two errors). The increase of overall mean RTs as a function of set size was highly reliable; the unconstrained model accounted for 99.81% of the variance of the overall mean RTs (Table 1). Slopes for *yes* and *no* responses were practically identical, with a slope ratio of 1.01 when freely estimated. When restricting the slopes to be equal, the slope for *yes* and *no* responses amounted to 55.2 ms/digit, and explained variance did not change (99.81%), implying that the slopes for *yes* and *no* responses were indeed equal. As both linear models explain practically all variance without quadratic terms, the increase of overall mean RTs as a function of set size can be taken to be linear (cf. Burrows & Okada, 1971; Chase & Calfee, 1969; Corbin & Marquer, 2008).

**Figure 2.**
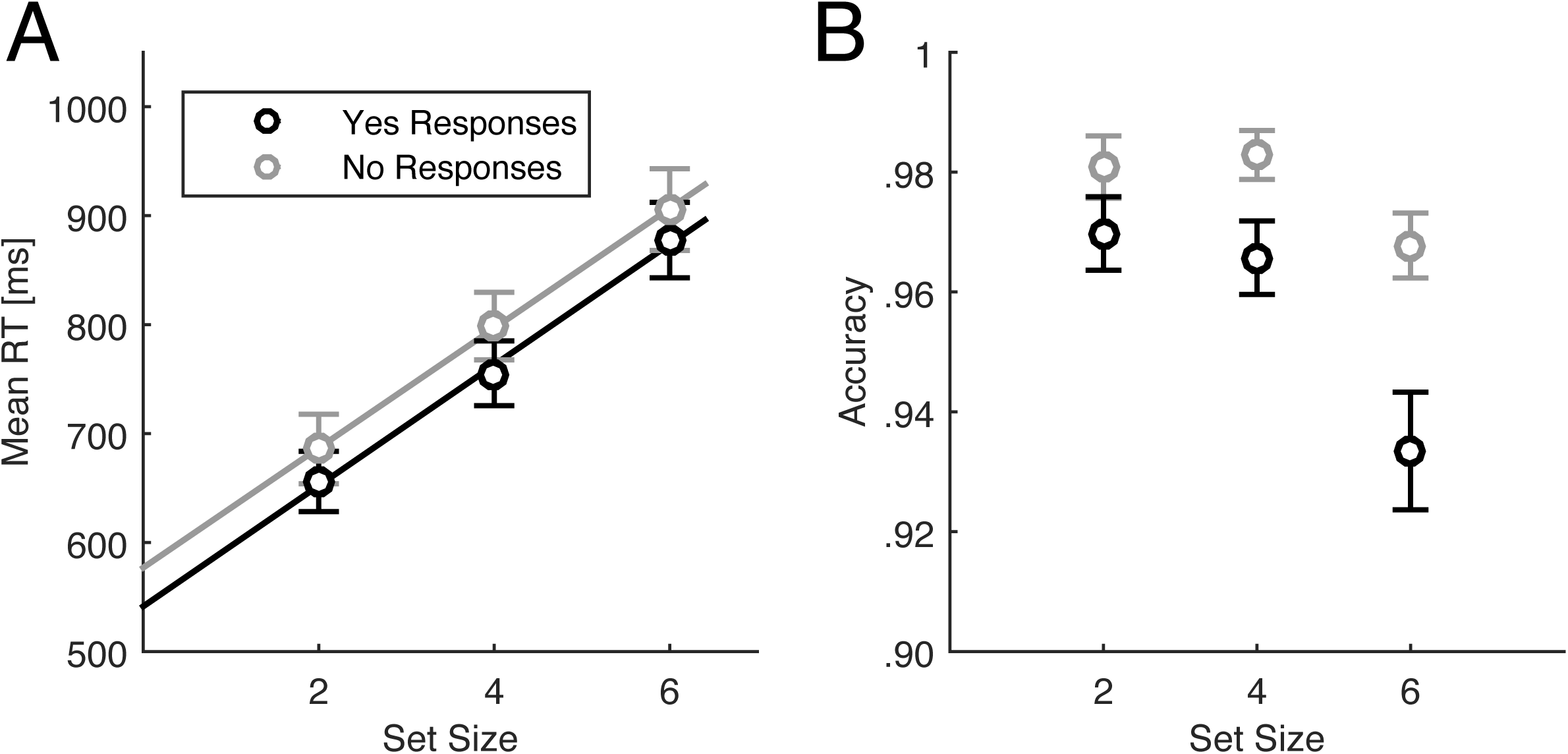
Overall mean response times (*A*) and accuracies (*B*) in session one. Slopes amounted 55.4 ± 2.4 (SE) ms/digit for *yes* responses and 54.9 ± 2.4 ms/digit for *no* responses. Slope ratio (positive slope divided by negative slope) was 1.01; positive and negative slopes did not differ reliably. Error bars indicate the standard errors of the mean.

**Table 1.** Comparison of parameters of the constrained and unconstrained linear model in session one.

|  | Unconstrained model |  |  | Constrained model |  |  |
| --- | --- | --- | --- | --- | --- | --- |
| | | $t(2)$ | $p$ | | $t(3)$ | $p$ |
| Intercept (+) [ms] | 541.4 | 51.62 | < .001 | 542.4 | 83.41 | < .001 |
| Intercept (−) [ms] | 577.0 | 55.01 | < .001 | 576.1 | 88.60 | < .001 |
| Slope (+) [ms/digit] | 55.4 | 22.82 | .002 | 55.2 | 39.19 | < .001 |
| Slope (−) [ms/digit] | 54.9 | 22.63 | .002 |  |  |  |
| $R^2$ [%] | 99.81 | | | 99.81 | | |

*Note.* In the constrained model, slopes are set to be equal. (+) = positive probes, (–) = negative probes.

Relying on MLM, we tested for equality of slopes more formally: Model fit did not change when constraining the slopes to be equal [Δ*χ*^2^(1) = 0.01, *p* = .943]. Furthermore, formal testing for linearity of slopes within MLM did not indicate reliable improvement of model fit when allowing the overall mean trend to be quadratic (Appendix B). Thus, for session one we conclude that slopes for *yes* and *no* responses are equal and linear.

It should be noted that intercepts of *yes* and *no* responses differed by 35.5 ms when slopes were freely estimated and by 33.7 ms when slopes were constrained to be equal for both response types, respectively. When adding the constraint of equal intercepts to the constrained model, MLM model fit reliably decreased [Δ*χ*^2^(1) = 12.61, *p* < .001]. Thus in contrast to the original report by Sternberg (1966), but in agreement with several other studies (e.g., Burrows & Okada, 1971; Corbin & Marquer, 2008, 2009; Nickerson, 1966; Ross, 1970), evidence for a difference in the intercepts was found in the current study.

#### Practice Effects

*Mean response times and accuracies.* A considerable—approximately one third— decrease of overall mean RTs as a function of practice was observed (Figure 3*A*). Omnibus rmANOVA indicated a reliable effect of repeated testing (sessions one to eight) on mean RTs; in addition, across sessions one to eight, effects of set size, probe type, as well as all two-way interactions were reliable (Table 2). Session by session comparisons of mean RTs within set sizes and probe types (paired-samples Student’s *t*-test, Figure 3*B*) indicated stable, asymptotic mean RTs on the group level from session six on. This was confirmed with post hoc rmANOVA on mean RTs from session six to eight, respectively (Table 2). No reliable change of overall mean RTs in sessions six to eight was found [*F*(1.87,58.07) = 0.28, *p* = .743]; the actual observed effect size of session amounted *f* = 0.09 (partial *η*^2^ = 0.01; average difference of mean RTs across sessions = 3.3 ms). Furthermore, no reliable interactions with the factor session were found. Thus, it can be concluded that at the group level asymptotic performance was achieved from session six on (mean level stability).

**Figure 3.**
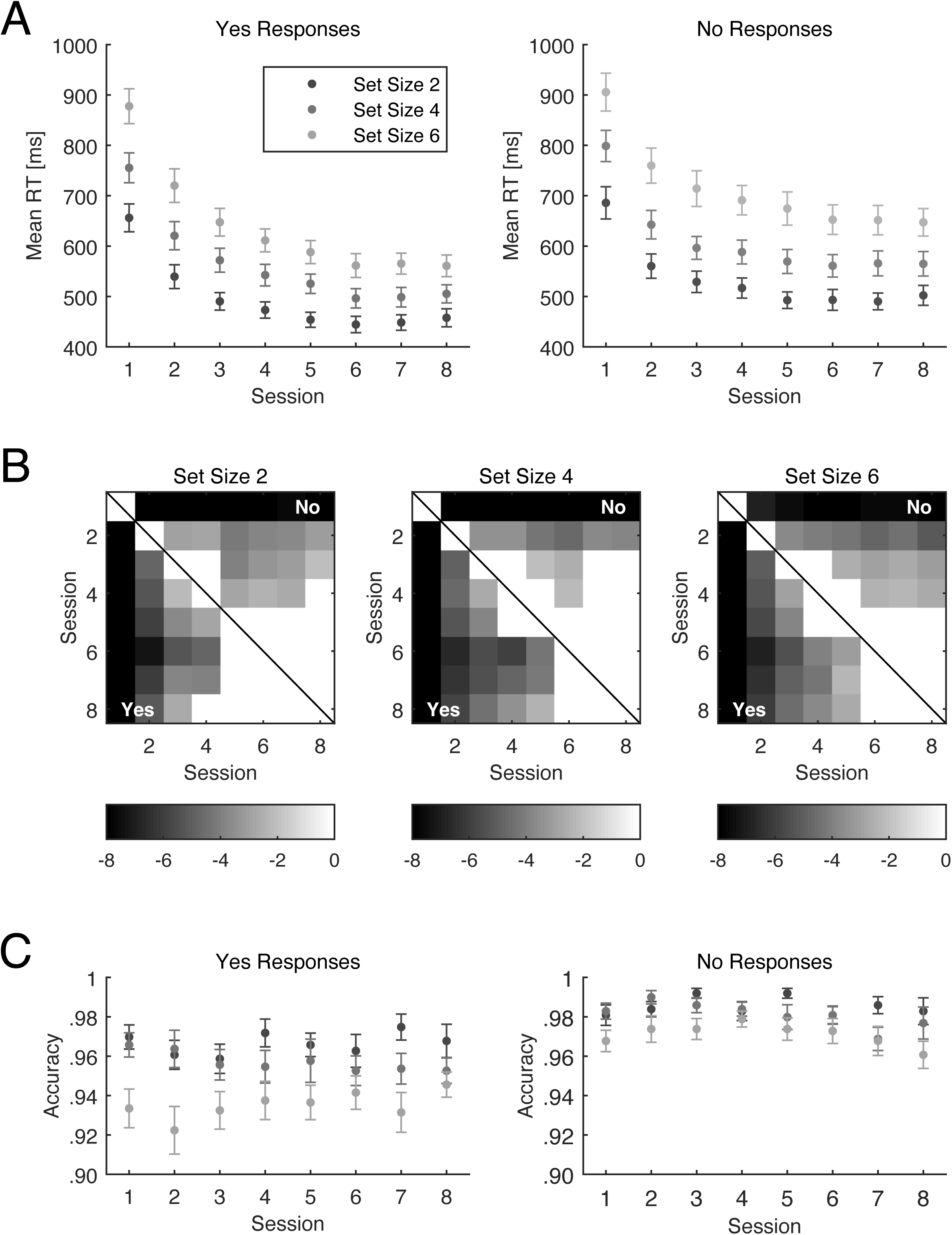
*A* Change in overall mean response times as a function of practice. *B* Pair-wise comparisons of mean response times within set size and response type across all sessions (paired-samples Student’s *t*-tests). *t* values are color coded (color bars indicate scaling of *t* values); non-significant (*α* level of .05) *t* values are masked out (white boxes). Lower triangles show comparisons of *yes* responses, upper triangles of *no* responses. From session six on no reliable change of overall mean response times was observed. Note that not correcting for multiple comparisons leads to a more conservative evaluation of achievement of asymptotic performance. *C* Accuracies across eight sessions; no apparent systematic change due to practice is discernible. Color coding same as in *A*. Error bars indicate the standard errors of the mean.

**Table 2.** rmANOVA results on mean RTs.

|  | Sessions 1–8 |  |  |  | Sessions 6–8 |  |  |  |
| --- | --- | --- | --- | --- | --- | --- | --- | --- |
| | $df_n$ | $df_d$ | $F$ | $p$ | $df_n$ | $df_d$ | $F$ | $p$ |
| A: Session | 1.85 | 57.45 | **75.67** | **< .001** | 1.87 | 58.07 | 0.28 | .743 |
| B: Set Size | 1.16 | 36.04 | **171.90** | **< .001** | 1.26 | 39.01 | **124.46** | **< .001** |
| C: Probe Type | 1.00 | 31.00 | **53.40** | **< .001** | 1.00 | 31.00 | **63.11** | **< .001** |
| A × B | 5.25 | 162.64 | **8.74** | **< .001** | 2.97 | 91.95 | 1.54 | .209 |
| A × C | 3.89 | 120.66 | **5.40** | **.001** | 1.94 | 60.00 | 0.35 | .700 |
| B × C | 1.18 | 36.63 | **6.56** | **.011** | 1.31 | 40.52 | **9.85** | **.001** |
| A × B × C | 7.40 | 229.46 | 1.45 | .182 | 3.51 | 108.73 | 0.19 | .929 |

*Note.* Degrees of freedom ( $df$ ) Greenhouse-Geisser corrected.

Overall mean accuracies from sessions one to eight did not show any systematic change over time (Figure 3*C*) and closely resembled the accuracies found in session one. rmANOVA did not reveal reliable effects of session [*F*(5.03,156.00) = 0.28, *p* = .923], and importantly, no interaction including the factor session [*F*s < 1.57, *p* > .163]. Thus, the decrease of overall mean RTs cannot be accounted for by a speed-accuracy trade-off.

*Intercepts.* Intercepts of the unconstrained model fitted to the mean RTs decreased with practice (Figure 4*A*), reflecting the pattern found for the mean RTs. At asymptote, *yes* response intercepts were found to range between 383.7 and 404.8 ms and *no* response intercepts were found to range between 407.8 and 426.4 ms (Table 3); analogous to session one, a systematic difference of 21.6 to 25.8 ms between *yes* and *no* response intercepts was observed.

**Figure 4.**
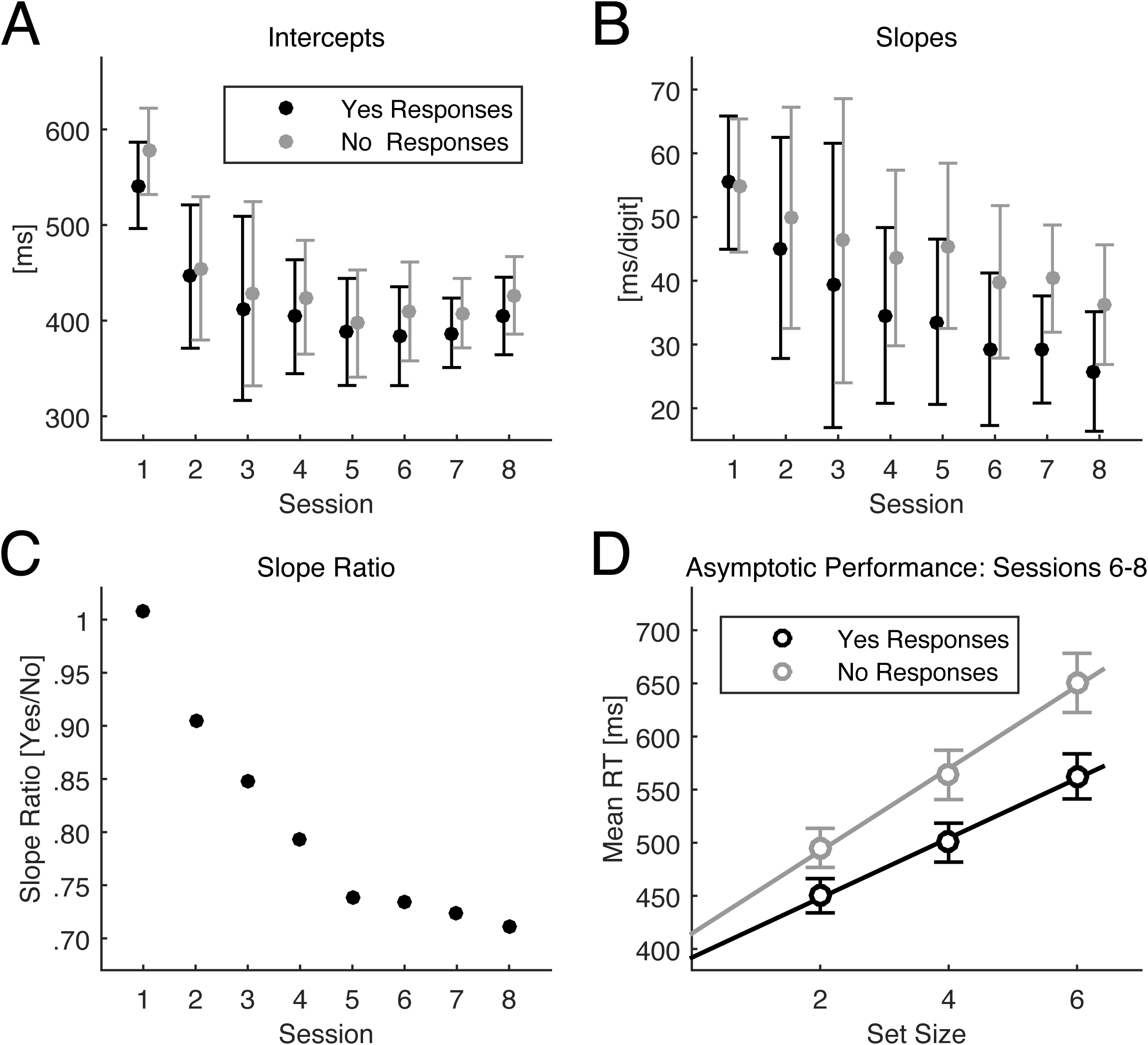
Intercepts (*A*) and slopes (*B*) of *yes* and *no* responses in the unconstrained model. Error bars indicate 95% confidence intervals of parameter estimates. *C* Slope ratio, *yes* divided by *no* response slopes. *D* Overall mean response times at asymptotic performance (sessions 6–8); slope ratio amounted 0.72. Error bars indicate standard errors of the mean.

**Table 3.**
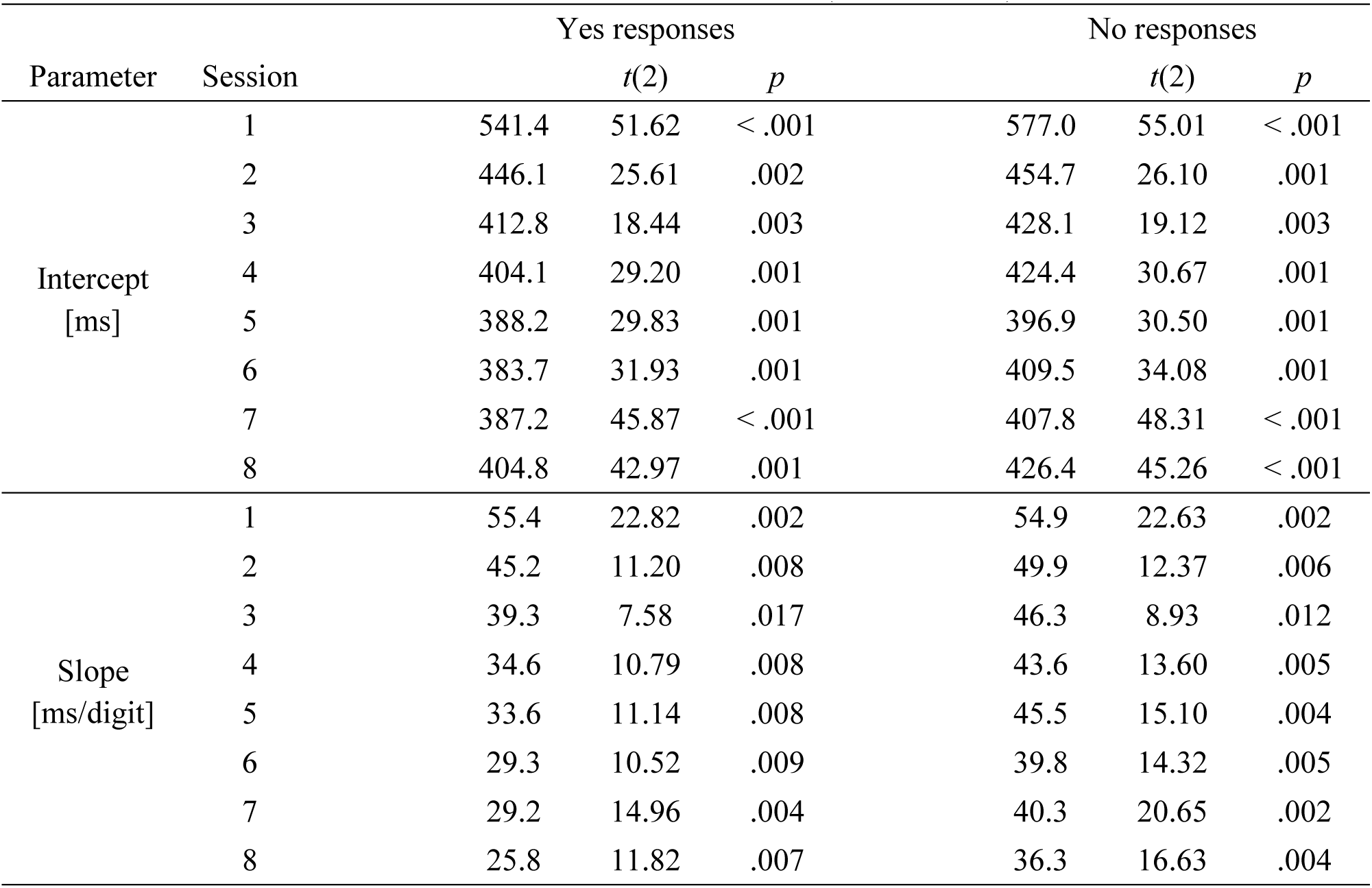
Parameter estimates for the unconstrained model (sessions 1–8).

*Slopes. Yes* and *no* response slopes decreased with practice (Figure 4*B*). However, the statistically reliable increase of RTs as a function of set size did not disappear with practice, and continued to be present when asymptotic performance was reached (Table 3; highly reliable set size effect in the rmANOVA for sessions 6–8, Table 2). At asymptote (sessions six to eight), mean *yes* response slopes were found to range between 25.8 and 29.3 ms/digit and mean *no* response slopes were found to range between 36.3 and 40.3 ms/digit. The very low unexplained (residual) variances (*R*^2^ > 99.3%, with the exception of session three; Table 4) suggest that linear slopes provide a viable account of the average data, especially with the unconstrained model (formal testing of linearity of slopes within MLM is provided in Appendix B). Thus, overall slopes were found to be reliably different from zero and remained linear after reaching asymptotic performance.

**Table 4.** Model fit statistics for and comparison of the unconstrained (A) and constrained (B, equal slopes) model across eight practice sessions.

| Session | Model fit indices |  |  |  | Model comparisons |  |
| --- | --- | --- | --- | --- | --- | --- |
| | 100-$R^2$ [%] | | -2 LL | | Model A vs. B | |
| | A | B | A | B | $\Delta\chi^2(1)$ | $p$ |
| 1 | 0.19 | 0.19 | 2224.43 | 2224.44 | 0.01 | .943 |
| 2 | 0.69 | 0.93 | 2151.80 | 2152.91 | 1.11 | .293 |
| 3 | 1.31 | 1.91 | 2139.90 | 2141.85 | 1.95 | .163 |
| 4 | 0.55 | 1.65 | 2115.01 | 2118.79 | 3.78 | .052 |
| 5 | 0.48 | 2.34 | 2062.91 | 2069.82 | **6.91** | **.009** |
| 6 | 0.46 | 2.14 | 2076.44 | 2082.90 | **6.47** | **.011** |
| 7 | 0.23 | 2.12 | 2014.60 | 2025.76 | **11.16** | **.001** |
| 8 | 0.35 | 2.35 | 2041.39 | 2047.43 | **6.04** | **.014** |

*Note.* Reliable differences in model fit (model comparison) are printed bold.

*Change of the mean slope ratio as a function of training.* The difference between slopes of *yes* and *no* responses progressively increased with practice (Figure 4*C*). Average slope ratios in sessions six to eight ranged from 0.71 to 0.74, also showing stabilization at asymptotic performance. These values are approximately half-way between the values predicted by the serial exhaustive (ratio = 1.0) and serial self-terminating (ratio = 0.5) memory search models. When testing for slope differences within MLM, reliable slope differences were found from session five onwards (Table 4). The overall mean RTs pooled across sessions six to eight (i.e., at asymptotic performance) as well as the unconstrained model fit are shown in Figure 4*D*.

Taken together, the model that constrains the slopes of *yes* and *no* responses to be equal provided an acceptable representation of the data in session one but proved to be an increasingly inadequate representation of the overall mean RT pattern with increasing amounts of practice (Table 4, Figure 4). At asymptotic performance, the unconstrained model fit the data reliably better than the constrained model, indicating reliable differences between the slopes of *yes* and *no* responses after a few days of practice at the group level.

### Towards the Individual Level

As discussed in the Introduction, investigating psychological processes at the individual level requires high reliability of data, which in turn is dependent on high mean and relative stability of data. In the following sections, we report our investigation of reliability and stability of mean RTs and model parameters in the Sternberg paradigm, and demonstrate that the reliability at the level of a single session was insufficient to make sound inferences about RT patterns at the individual level. Hence, we investigated whether reliability increased as predicted by the Spearman-Brown formula when pooling data across multiple sessions, and retraced its dependence on the stability of data.

#### Within Session Split-half Reliability

Split-half reliability of mean RTs was found to be high and homogeneous (*ρ* = .94 – .96; Figure 5). Split-half reliability of intercepts was also found to be high but systematically lower (*ρ* = .79 – .88), and split-half reliability of slopes was found to be even lower (*ρ* = .71 – .81). Finally, split-half reliability of the slope ratio was rather low (*ρ* = .31 – .55) and on a level practically impeding attribution or identification of a cognitive process at the individual level. Across sessions, reliability of mean RTs as well as model parameters did not change systematically; thus, practice per se did not influence the reliability of data and parameters.

**Figure 5.**
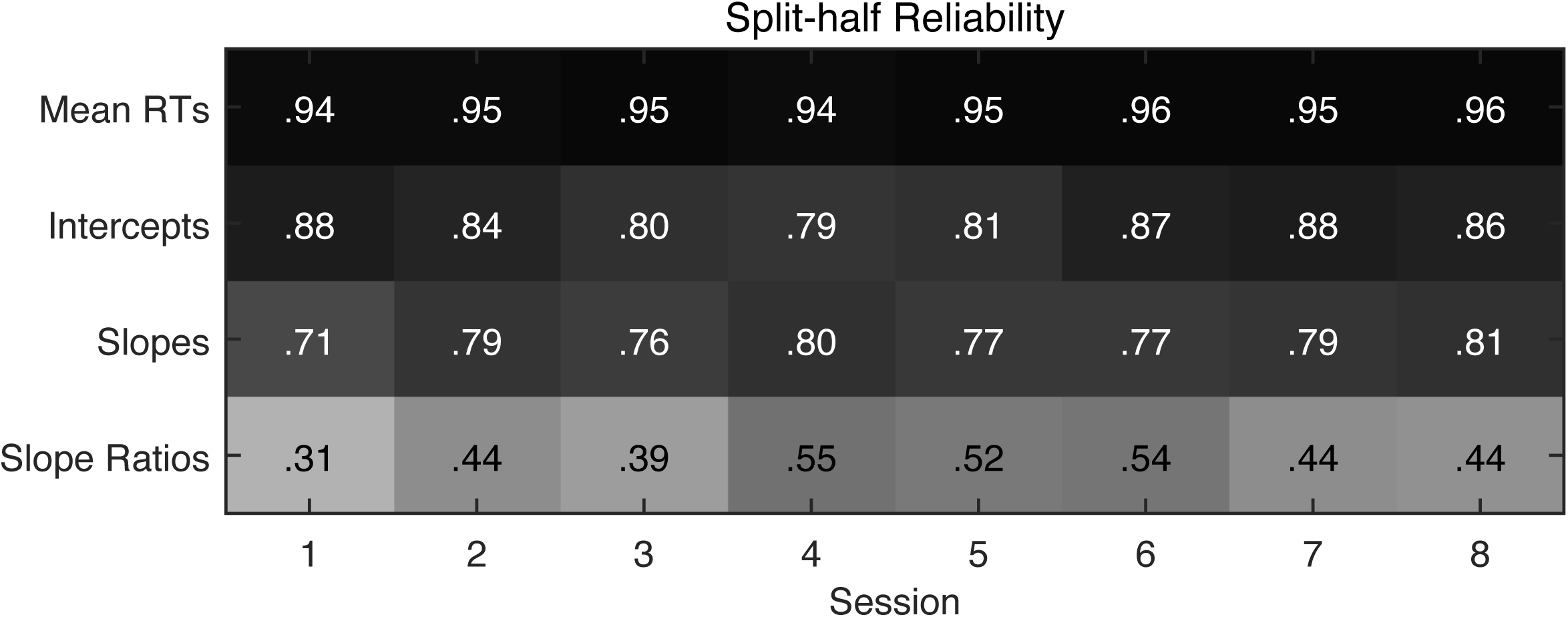
Average split-half reliabilities of mean response times (RTs), intercepts, slopes, and split-half reliabilities of slope ratios across eight sessions.

#### Relative Stability

In addition to split-half reliability, we assessed the relative (i.e., retest) stability of mean RTs and model parameters across sessions. For mean RTs, a medium to high level of stability was observed (see Figure 6; *ρ* = .67 – .93). Noticeably, relative stability near asymptotic performance (i.e., in sessions five/six to eight) was found to be on a homogeneously high level, in contrast to stability during the initial practice sessions. Furthermore, relative stability between more distant sessions was generally lower than relative stability between adjacent sessions. Statistical comparisons between the average stability coefficients of the first four versus the latter four sessions as well as stability coefficients of ‘distant’ sessions (i.e., between sessions separated by at least three sessions in between) are provided in Table 5. The average stability coefficient of sessions five to eight were significantly higher than the average stability coefficients of sessions one to four and of the distant sessions.

**Figure 6.**
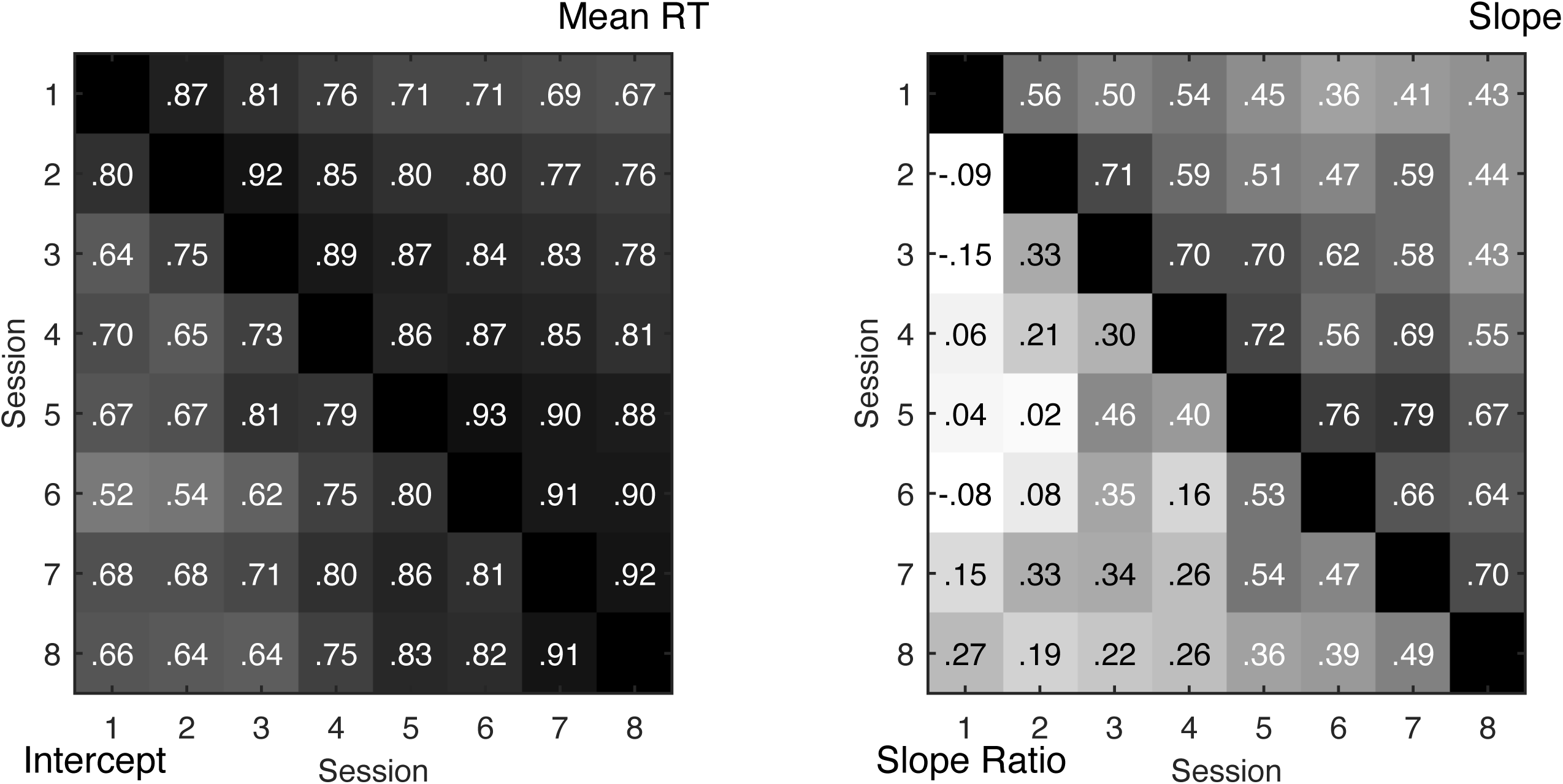
Relative stability coefficients of mean RTs, intercepts (left upper and lower triangle), slopes, and slope ratios (right upper and lower triangle).

**Table 5.**
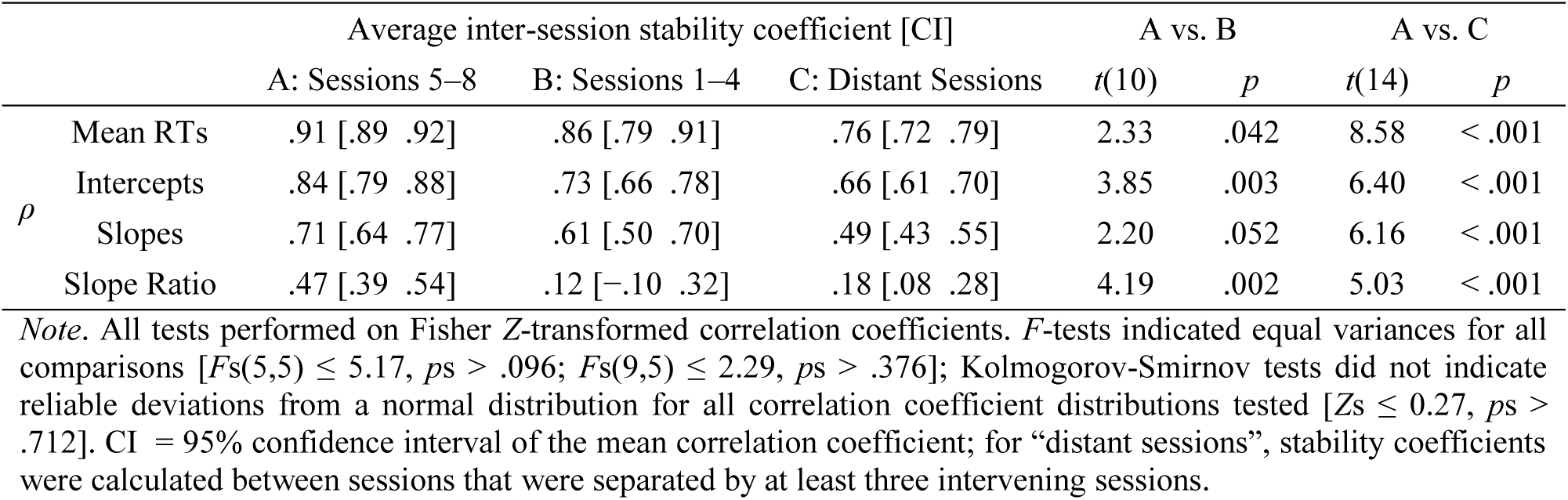
Comparison of average inter-session stability coefficients at different stages of the practice phase.

The general pattern of relative stability of the model parameters (Figure 6) followed the pattern described above for the mean RTs. Furthermore, in parallel to the split-half reliability, overall stability decreased systematically from mean RTs, to intercepts, slopes, and slope ratios. For intercepts, the stability was predominantly medium to high (*ρ* = .52 – .91), for slopes the stability was found to be systematically lower (*ρ* = .36 – .79), and for the slope ratio, stability was found to be distinctly low, including zero (*ρ* = −.14 – .55). Overall, the average relative stability was found to be reliably higher near asymptote than during the initial practice sessions and across distant sessions (Table 5).

Taken together, relative stability was lower during the first four sessions, indicating changes in the rank order of participants that are likely to reflect individual differences in practice gains. Consequently, initial performance was not entirely predictive of asymptotic performance. The most important indicator for the identification of memory scanning processes—the slope ratio—showed a level of relative stability across the initial sessions that rules out sensible identification of person parameters based on typical single-session data. Higher levels of relative stability were observed near asymptotic performance, that is, coinciding with high levels of mean stability.

#### Split-half Reliability when Pooling Data across Sessions

Given that reliability of the slope ratio was too low for identifying memory scanning processes at the individual level, we explored whether pooling data across sessions would increase the reliability of model parameters as predicted by the Spearman-Brown formula.

The comparison between observed average split-half reliabilities of mean RTs, intercepts, slopes, and slope ratios and the corresponding coefficients predicted by the Spearman-Brown formula based on the average single session split-half reliabilities are provided in Table 6 and Figure 7. Descriptively, pooling of data from sessions six to eight lead to a consistent increase of observed split-half reliabilities for all reported measures; in contrast, pooling of data from sessions one to three did not lead to an equally consistent increase. The divergence between predicted and observed coefficients was quantified by testing the equality of the observed correlation coefficients with the predicted correlation coefficients; resulting *z*-values are reported in Table 6. Observed and predicted split-half reliabilities are more similar to each other in sessions six to eight than in sessions one to three. When pooling data from three sessions, divergence from the predicted coefficients was | *z* | = 0.02 – 1.02 for sessions six to eight as compared to | z | = 1.26 – 2.20 for sessions one to three. For the slope ratio, observed split-half reliabilities of pooled sessions one to three (*ρ* = .35) and six to eight (*ρ* = .73) were found to be reliably different (Fisher’s *z*-test: | *z* | = 2.17, *p* = .030).

**Figure 7.**
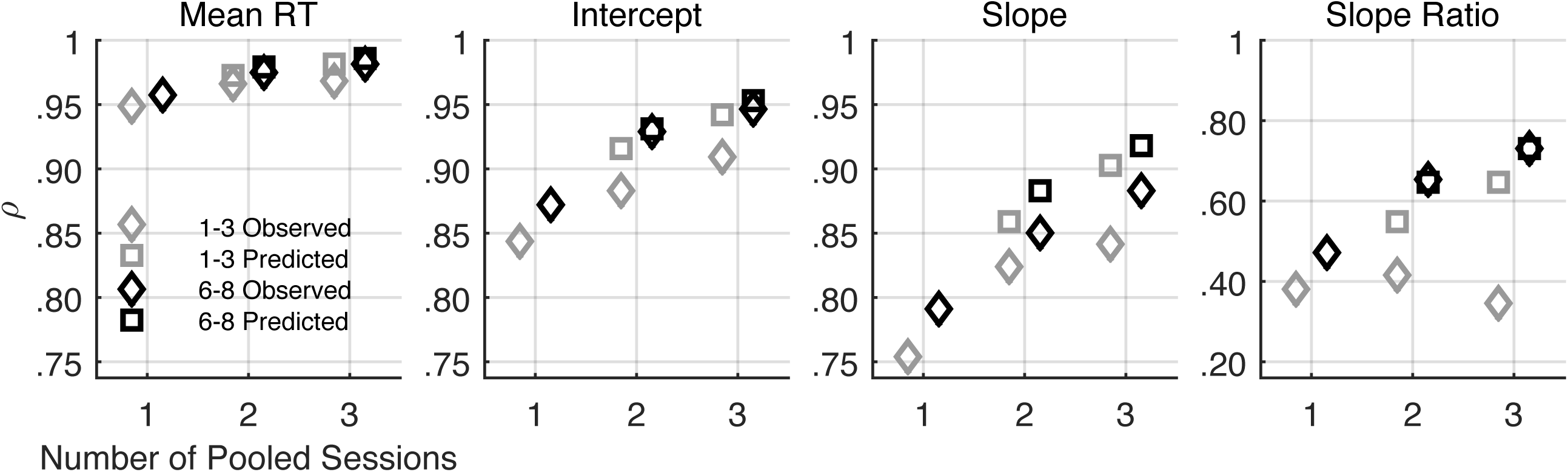
Comparison of predicted and observed split-half reliabilities (*ρ*) when increasing the amount of data by pooling data across up to three sessions. Note the change of the scale of the y-axis for the slope ratio. Overall, observed reliabilities follow the predicted values more closely when aggregating data from sessions six to eight. 1–3 = pooling of sessions one to three; 6–8 = pooling of sessions six to eight.

**Table 6.** Comparison of observed and predicted split-half reliabilities (*ρ*) when aggregating data across two and three sessions.

| Sessions | # Ses Pooled | Mean RTs |  |  | Intercepts |  |  | Slopes |  |  | Slope Ratio |  |  |
| --- | --- | --- | --- | --- | --- | --- | --- | --- | --- | --- | --- | --- | --- |
| | | Obs | Pred | $z$ | Obs | Pred | $z$ | Obs | Pred | $z$ | Obs | Pred | $z$ |
| 1–3 | 1 | .95 |  |  | .84 |  |  | .75 |  |  | .38 |  |  |
|  | 2 | .97 | .97 | –0.78 | .88 | .92 | –0.93 | .82 | .86 | –0.66 | .42 | .55 | –0.95 |
|  | 3 | .97 | .98 | –1.67 | .91 | .94 | –1.26 | .84 | .90 | –1.39 | .35 | .65 | –2.20 |
| 6–8 | 1 | .96 |  |  | .87 |  |  | .79 |  |  | .47 |  |  |
|  | 2 | .97 | .98 | –0.48 | .93 | .93 | –0.10 | .85 | .88 | –0.72 | .65 | .64 | 0.06 |
|  | 3 | .98 | .99 | –0.76 | .95 | .95 | –0.41 | .88 | .92 | –1.02 | .73 | .73 | 0.02 |

*Note.* Pooling across sessions is systematically less advantageous for sessions one to three. # Ses Pooled = number of sessions pooled; Obs = observed reliability; Pred = reliability predicted by the Spearman-Brown formula from average single session split-half reliability; $z$ = “effect size” of difference between predicted and observed correlation coefficient.

Taken together, pooling three sessions at asymptotic performance systematically increased reliability and was more beneficial to reliability than pooling sessions in the presence of strong practice effects; this was especially true for the parameter of highest interest, the slope ratio.

### Individual Level

After having established the achievement of asymptotic performance within six sessions, with high mean as well as relative stability, we proceed with reporting individual differences observed at asymptote. By pooling sessions six to eight, we capitalize on the improved reliability in order to investigate the identifiability of memory search processes at the individual level, and contrast these with the single session data from session one for illustrative purposes.

Figure 8 summarizes single participants’ mean RTs aggregated across sessions six to eight together with the least-squares regression lines of the unconstrained model. It can be seen that overall fits are satisfying (*R^2^* = 90.9% to 100%). Figure 9 provides bootstrapped 95% confidence intervals (CIs) for the slope ratios of all 32 participants. 13 (40.6%) participants’ CIs contain a slope ratio of 0.5, denoting self-terminating memory search, and 13 participants’ CIs contain a slope ratio of 1.0, denoting exhaustive memory search. Thus, 26 of 32 participants (81.2%) provided data that were consistent with one of the two competing memory search models, with the same number of participants falling into either category. Of the remaining participants, one (3.1%) had a slope ratio reliably below 0.5, three (9.4%) had slope ratios reliably above 1.0, and two (6.2%) participants’ CIs included slope ratios of 0.5 and 1.0.

**Figure 8.**
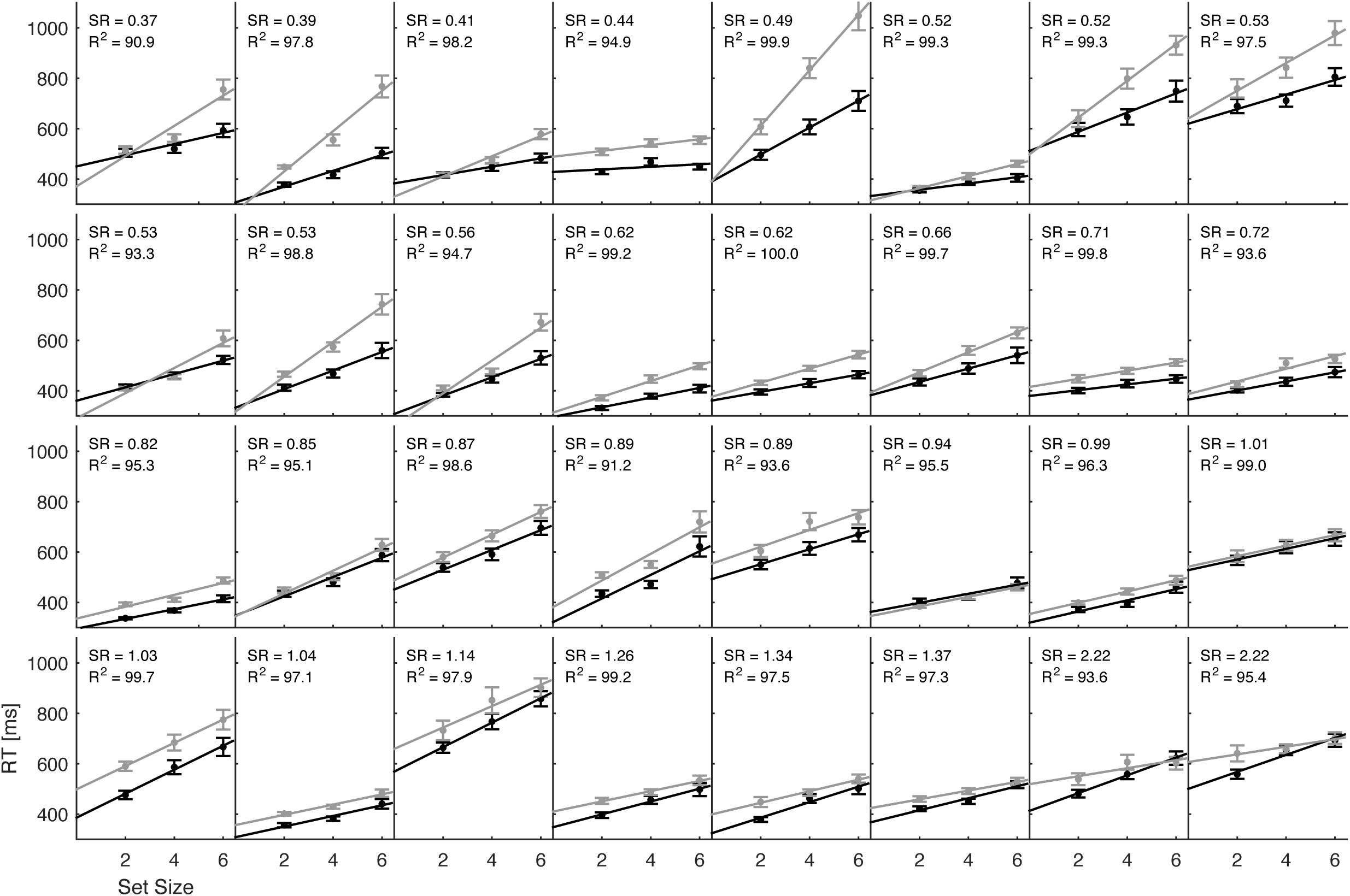
Mean response times and least-squares regression lines for individual participants at asymptotic performance (sessions six to eight); black = *yes* responses, gray = *no* responses. SR = slope ratio. Error bars indicate the standard error of the mean.

**Figure 9.**
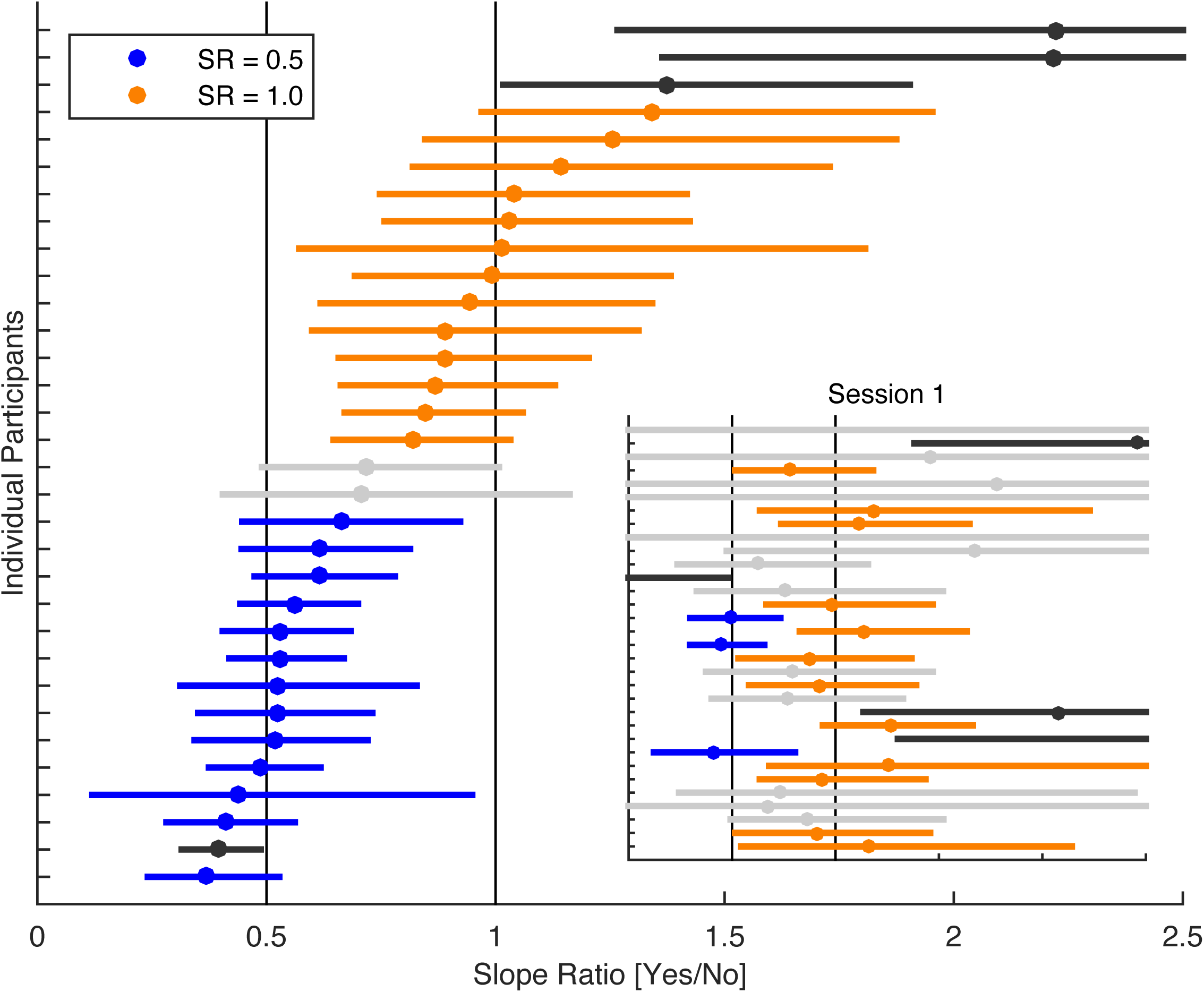
Individual participants’ bootstrapped 95% confidence intervals (CIs; horizontal lines) of slope ratios; data pooled across sessions six to eight. 26 participants’ CIs contained either slope ratio 0.5 or 1, denoting self-terminating (slope ratio = 0.5) or exhaustive (slope ratio = 1) STM search, respectively. Two participants’ CIs contained both slope ratios (light gray), four participants’ CIS contained neither slope ratio (dark gray). There are considerable individual differences in reliability of slope ratios, as expressed by the differing individual CIs. Inset shows slope ratios and 95% CIs from session one; the order of participants corresponds to the order of participants in sessions six to eight. SR = slope ratio.

Finite mixture modeling using Gaussian mixture distributions favored a three component solution (AIC = 13.9) over two or one component solutions (AIC ≥ 16.5). The mean parameters in the three component model were estimated to be 0.51, 0.94, and 2.22, with a relative proportion of 37.4%, 56.4%, and 6.2%, respectively. Thus, the majority of slope ratios were captured by two Gaussian distributions centered almost exactly at the expected slope ratios of the two competing memory search models.

As can be seen from this perspective, the superiority in fit of the unconstrained over the constrained model (cf. Table 4, Figure 4) reflects the presence of a large proportion of participants with slope ratios reliably lower than 1.0. At the same time, it is worth noting that the best-fitting model for the overall mean responses (Figure 4*D*) does not capture the full range of slope ratios observed at the individual level. The CI of only 16 (50%) participants comprised the slope ratio of 0.72 obtained at the group level for sessions six to eight. In other words, for half the participants, the overall mean model cannot be rejected, but for the other half of the participants the mean model does not present a viable account of their RT pattern.

The inset in Figure 9 provides the slope ratios and CIs of individual participants from session one. CIs are considerably larger as compared to the pooled data at asymptotic performance; the CIs of 13 (40.6%) individuals contained the slope ratio of both 0.5 and 1.0; the CIs of four (12.5%) individuals contained neither. For three (9.4%) participants, the slope ratio of 0.5 was included in their CI, and for twelve (37.5%) individuals, the slope ratio of 1.0 was included in their CIs. Thus, for over one third (40.6%) of the participants, attribution of one of the two alternative search models was not possible in the first session. In correspondence with the overall mean model, which was favoring the exhaustive search model in session one, the majority of identifiable slope ratios did not differ reliably from 1.0. Nonetheless, even at the single session level, some individuals defy the generality of the exhaustive search model. Furthermore, the association between the slope ratios at asymptotic performance and the slope ratios observed in the first session did not differ reliably from zero, *ρ* = .07 (*p* = .670).

Taken together, pooling data at asymptotic performance with high mean as well as relative stability increased the identifiability of memory search processes at the individual level and revealed reliable and substantial individual differences. The majority of participants provided data consistent with one of the two major competing memory search models, serial exhaustive and self-terminating memory search, thus, prohibiting the adequate description of memory search with a single model; the model fitted that fitted the overall mean data convincingly was not generalizable to the individual level.

### Differences between ‘Exhaustive’ and ‘Self-terminating’ Memory Searchers

As the majority of participants could be subsumed in two groups, we examined whether individuals complying with either the exhaustive (*n* = 13) or self-terminating (*n* = 13) memory search model at asymptotic performance differed in the assessed covariates and parameters of the Sternberg paradigm (Table 7). In the covariates, groups did not differ reliably from an equal distribution of female and male participants, as well as in chronological age, number of school years, general fluid intelligence (Raven Matrices), and all assessed memory performance scores with the exception of Reverse Span (*d* = 1.00). In the parameters of the Sternberg task, a reliable group difference in the average *no* response slopes^1^ at asymptotic performance (*d* = −0.95) was found. For mean RTs, mean SDs, and accuracies no reliable differences were observed. Given the small sample size, balancing statistical power and type I error in a principled way was restricted. Validity of the observed differences may be gained from the following additional observations: (1) reliable effects are consistent (consistently better average performance in the exhaustive memory search group), (2) the group difference in the Reverse Span task is consistent in direction with all other memory span tasks assessed (Digit Span, WM Sorting, Operation and Symmetry Span), and (3) group differences in the *no* response slopes are consistent with the literature (cf. Sternberg, 1975). However, Figure 10 provides an important qualification of the group differences as the distributions of the groups largely overlap. The reverse span scores and *no* response slopes of the majority of the self-terminating group are within the range of the exhaustive group. Thus, with respect to the individual level, self-terminating search is more likely but not necessarily associated with lower memory span and slower memory scanning rates.

**Figure 10.**
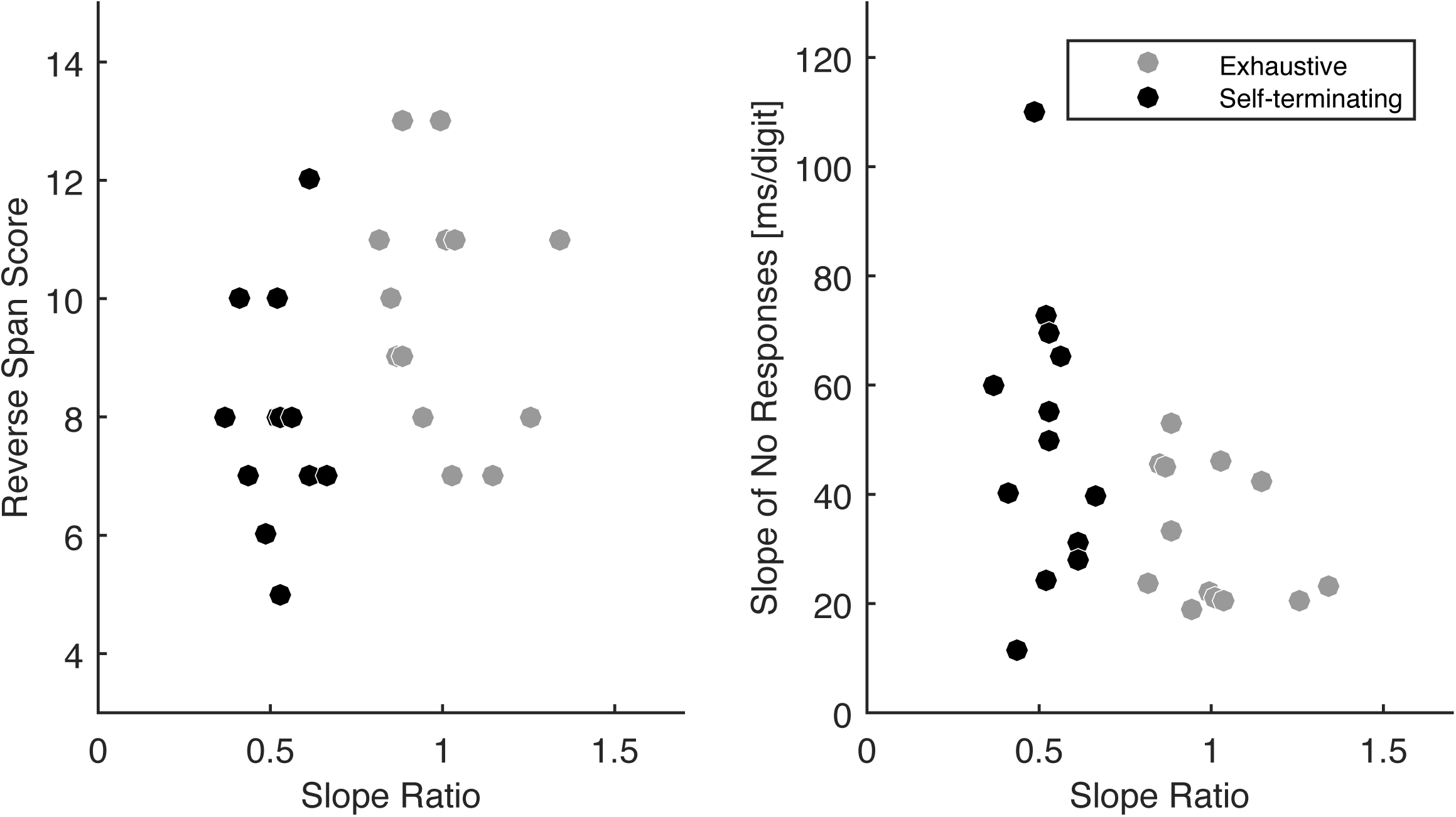
Comparison of the Reverse Span scores (covariate session) and slopes of *no* responses (sessions six to eight) of exhaustive (*n* = 13) and self-terminating (*n* = 13) memory searchers.

**Table 7.** Post hoc assessment of differences between individuals complying with the exhaustive or self-terminating memory search model at asymptotic performance.

| Task / Parameter |  | Exhaustive<br>(*n* = 13) | Self-terminating<br>(*n* = 13) | *t*(24) | *p* |
| --- | --- | --- | --- | --- | --- |
| Covariates | Female | 6 (46.2%) | 8 (61.5%)<sup>1</sup> |  |  |
|  | Age | 22.8 ± 1.6 | 23.0 ± 2.0 | −0.213 | .833 |
|  | School Years | 12.8 ± 0.6 | 12.3 ± 0.9 | 1.647 | .113 |
|  | University Students | 13 (100%) | 11 (84.5%) |  |  |
|  | Digit Span | 11.1 ± 2.9 | 9.9 ± 1.8 | 1.210 | .238 |
|  | Reverse Span | 9.9 ± 2.0 | 8.00 ± 1.8 | **2.435** | **.023** |
|  | WM Sorting | 10.7 ± 2.7 | 9.3 ± 2.5 | 1.352 | .189 |
|  | Operation Span I | 55.9 ± 14.6 | 54.6 ± 12.4 | 0.246 | .808 |
|  | Operation Span II | 56.2 ± 10.3 | 49.8 ± 12.6 | 1.416 | .170 |
|  | Symmetry Span | 33.2 ± 7.8 | 28.8 ± 6.6 | 1.553 | .134 |
|  | Digit Symbol | 70.6 ± 8.8 | 72.1 ± 11.3 | −0.368 | .716 |
|  | Raven Matrices | 11.1 ± 3.7 | 11.5 ± 3.5 | −0.274 | .786 |
| Sternberg task<br>Parameters | Mean RT [ms]<br>Session 1 | 770 ± 190 | 784 ± 160 | −0.198 | .845 |
|  | Mean RT [ms]<br>Sessions 6–8 | 539 ± 115 | 543 ± 122 | −0.086 | .932 |
|  | Mean SD [ms]<br>Session 1 | 170 ± 65 | 187 ± 59 | −0.688 | .498 |
|  | Mean SD [ms]<br>Sessions 6–8 | 93 ± 34 | 99 ± 43 | −0.348 | .731 |
|  | Mean Accuracy [%]<br>Session 1 | 97.2 ± 1.6 | 96.7 ± 2.0 | 0.617 | .543 |
|  | Mean Accuracy [%]<br>Sessions 6–8 | 96.0 ± 2.2 | 97.2 ± 2.2 | −1.269 | .217 |
|  | Slope [ms/item]<br>Session 1 | 49.2 ± 32.0 | 64.1 ± 32.6 | −1.127 | .271 |
|  | Slope [ms/item]<br>Sessions 6–8 | 31.9 ± 12.1 | 50.6 ± 24.9 | **−2.336** | **.028** |

*Note.* Slope = slope of *no* responses. Values are mean ± 1 *SD* across participants. Non-parametric Mann-Whitney *U*-tests revealed the same pattern of reliable results (reverse span: $z = -2.215$ , $p = .027$ ; *no* response slope in sessions 6-8: $z = -2.154$ , $p = .031$ ; all other comparisons $p > .05$ ). <sup>1</sup> $\chi^2(1) = .692$ , $p = .405$ when testing for equal distribution of females and males.

## Discussion

The rationale of this article was to explore in a principled way whether search from memory as assessed with the Sternberg paradigm is serial and exhaustive at the individual level. Empirical evidence in support of serial and exhaustive search from memory has been obtained with single-session group data. However, data gathered within a single session has been found to be insufficient to discriminate between serial exhaustive and alternative search processes at the individual level because the critical parameter, that is, the slope ratio, could not be estimated with sufficient reliability. Increasing data density by running multiple sessions permitted more reliable process identification at the individual level near asymptotic levels of performance. When slope ratios were estimated near asymptotic performance, memory search could be subsumed under exhaustive in 13 and self-terminating memory search in another 13 of 32 individuals. We conclude that individuals may differ reliably in the way they search information in memory and hence provide an empirical case where single session group data does not generalize to the behavior of all individuals. In the following, we retrace the main results of the study, and discuss their implications for cognitive psychology and neuroscience.

### Replication and Mean Level Perspective

In the first session, we replicated the core findings of Sternberg at the aggregate level (cf. 1966; 1969, 1975). This adds to our confidence that the implementation of the Sternberg task in this study did not deviate substantially from the original paradigm, and that the present sample as a whole did not differ in some unknown way from other convenience samples of healthy, younger adults.

Practicing the Sternberg task led to an overall reduction in mean RTs, intercepts, as well as slopes for *yes* and *no* responses. From session six onwards, mean RTs reached asymptotic levels. Importantly, even at asymptotic performance, slopes in the aggregate were linear and reliably different from zero. This indicates that fundamental characteristics (but not absolute parameter values) of the memory search process remained invariant with practice, which is an important prerequisite for relying on multiple repetitions of the task in order to increase data density without constraining the validity of measurements with respect to the cognitive process under investigation.

However, in contrast to the replication of the original Sternberg findings in session one, the data at asymptote were not well captured by the serial exhaustive model. At the aggregate level, slopes were found to differ reliably between *yes* and *no* responses from session five onwards, with a slope ratio halfway in between the slope ratios predicted from serial exhaustive and self-terminating search. Thus, at the mean level, the model from session one—comparable to a typical one-shot measurement—did not generalize to the (average) behavior near asymptotic performance. As will be discussed below, the analysis of individual data added an important qualification to this observation. Individuals differed reliably in their slope ratios at asymptotic performance, indicating that the mean model emanated from a mixture of individuals whose search was exhaustive and individuals whose search was self-terminating (Figure 9).

Also with respect to the generality of the memory scanning rate for digits of approximately 40 ms/digit as suggested by Sternberg (1966, 1969, 1975) a differentiated picture emerged. In line with other reports, we observed an average slope of approximately 55 ms/digit in session one (cf. Corbin & Marquer, 2008, 2009) and a decrease in the slopes as a consequence of practice to approximately 40 ms/digit at asymptotic performance (cf. Carter et al., 1980). In addition, substantial and reliable individual differences in the slopes were observed, well in line with the literature (cf. H. L. Brown & Kirsner, 1980; Carter et al., 1980; Carter et al., 1986; Chiang & Atkinson, 1976; Longstreth & Madigan, 1982; Puckett & Kausler, 1984; Roznowski & Smith, 1993). Thus, also the average slope from session one did not generalize to practiced (and individual) performance.

Taken together, these results show that the cognitive processes that are active during memory search may quantitatively and qualitatively change within (some) individuals over time (as implied by the group data). In line with our findings, it has been argued that memory search processes may be better discernible after some practice when processing reliably stabilizes at an efficient search strategy (Houck & Hoffman, 1986, after van Zandt and Townsend, 1993). Furthermore, from a conceptual point of view, one may tacitly assume that a cognitive process identified in the laboratory is similarly used in everyday life (but see for example Dhami, Hertwig, & Hoffrage, 2004). To the extent that everyday cognitive functioning relies on overlearned processes, it is plausible to argue that cognitive processes with relevance to ‘everyday’ cognitive functioning should be assessed in some practiced state. The attainment of stable performance levels after a few days of practice suggests that healthy young adults can quickly adapt to specific task requirements, likely by optimizing existing cognitive (and neural) processes within their currently available resources (cf. Clark, Appelbaum, van den Berg, Mitroff, & Woldorff, 2015; Lövdén et al., 2010). In sum, these observations also raise the interesting question, to what extent findings in experimental (as well as differential psychology) will provide a more differentiated picture when cognitive processes (and individual differences) are assessed in a practiced state (cf. Ackerman, 1988; M. M. Baltes, Kuhl, & Sowarka, 1992; Rogers et al., 2000), and how this influences their qualification as ‘general’.

### Approaching the Individual Level

The reliability of the slope ratios in the first session was insufficient to unequivocally identify cognitive processes at the individual level and to explore whether exhaustive search was present in all participants in this session (Figure 5; Figure 9, inset; see also H. L. Brown & Kirsner, 1980; Carter et al., 1980; Carter et al., 1986; Chiang & Atkinson, 1976; Longstreth & Madigan, 1982). Thus, we investigated whether reliability increased as predicted by the Spearman-Brown formula when pooling data across multiple sessions and evaluated the dependency of reliability on high mean and relative stability of the data across sessions.

At the single session level, the reliabilities of mean RTs were high (*ρ* ≥ .94; Figure 5). Mean RTs are observed scores that reflect a mixture of different processes underlying the execution of a cognitive task, and hence are less interpretable than the model parameters (e.g., slopes or slope ratios). Nevertheless, it is instructive to consider the extent to which model parameters differ in reliability from these observed scores. Notably, in contrast to the exceptional high reliability of the mean RTs, reliabilities for slope ratios were so low that identification of systematic differences or commonalities across individuals was close to impossible (*ρ* = .31 – .55; Figure 9, inset).

Relative (i.e., rank order) stability of mean RTs and parameters across multiple sessions increased with practice and apparently plateaued in the sessions in which asymptotic performance at the group level had been attained (Figure 6, Table 5). Near asymptote, relative stability was close to the split-half reliability (within sessions), which implies that the observed instability was well accounted for by unreliability. Furthermore, the high levels of relative stability in these later sessions indicate that the asymptotic nature of performance at the aggregate level is likely to generalize to the individual. Thus, from this perspective, the requirements for data aggregation within individuals with the purpose of noise reduction—stable performance level and variance attributable to random error—appear to be well met at asymptotic performance. Furthermore, the achievement of high relative stability in addition to the high mean stability after limited practice supports the above stated assertion that healthy younger adults are able to adapt to specific task requirements with limited amounts of practice.

Given the low reliability of the slope ratio for individual sessions, we explored whether pooling the RTs from multiple sessions would increase reliability as predicted by the Spearman-Brown formula. These explorations led to two main results. First, reliability increased by pooling data across up to three sessions (Figure 7). Second, observed reliabilities were systematically closer to the predicted reliabilities when pooling data near asymptotic performance than when pooling data from initial sessions (Table 6). The reliability of the slope ratio increased up to *ρ* = .73 when aggregating data from sessions six to eight, but it did not increase at all when pooling data from the first three sessions (*ρ* = .35). Thus, for the parameter of highest interest, the increments in reliability predicted by the Spearman-Brown formula were restricted to sessions in which performance had stabilized, well in line with theoretical considerations.

Presumably, an unknown but major portion of the early fluctuations in performance did not qualify as random noise but may indicate differential practice effects or the participants’ exploration of the task space (cf. Li, Huxhold, & Schmiedek, 2004). This is supported by the observation that the discrepancy between relative stability and split-half reliability was slightly larger in the initial four sessions, whereas the split-half reliability did not change systematically across sessions (i.e., practice per se did not influence the reliability). When aggregating data across sessions, the differential change of individual performance is apportioned to the unexplained variance, thus impairing reliability. Taken together, these observations underscore the importance of measuring individual behavior after the cognitive process of interest has stabilized, when repeated assessments are required in order to improve reliability.

### Individual Level

As noted above, the memory search process was not unequivocally identifiable at the individual level in the first session (Figure 9, inset). Nevertheless, despite the group data were well captured by the serial exhaustive memory search model, the picture at the individual level flagged a non-negligible degree of ambiguity and heterogeneity in the memory search process, as three individuals appeared to reliably engage in self-terminating search. When increasing the number of data points by aggregating three sessions at asymptotic performance, process identification at the individual level improved considerably. The majority of participants could be unambiguously assigned either to an exhaustive (40.6%) or a self-terminating (40.6%) memory search process. This observation also helps to explain why the slope ratio for the group (0.72) fell half way in between the two alternative memory search models (Figure 4*D*).

A remarkable and non-trivial finding of our study is that the search process seemingly used in the initial session does not predict the search process participants settle in after five days of practicing the task, as indicated by the lack of an association between the slope ratios observed in session one and the slope ratios near asymptote (*ρ* = .07). This finding may in part reflect the lack of reliability of the slope ratios in session one. However, for some individuals who already showed narrow CIs in session one, the search process had differed markedly near the asymptote (e.g., 10^th^ and 18^th^ individual from the bottom in Figure 9). Together with the failure to increase reliability in the early part of the experiment by pooling data across sessions, this observation adds to the impression that a fair number of individuals used more than one search process in the early period of the experiment, and settled into a more stable state of either predominantly self-terminating or exhaustive search later on (cf. Van Zandt & Townsend, 1993).

In line with this conjecture, participants exhibited substantial differences in the reliability of their slope ratios, as indicated by their CIs (Figure 9). Even near asymptotes, the point measures of the slope ratios are not equally trustworthy across participants. Data quality may differ across participants, for example due to differences or fluctuations in attention and motivation. In addition, it is conceivable that some participants continue to apply different cognitive processes across individual trials. In this case the width of the CIs may also reflect differential strategy use within individuals, in line with findings reported by Corbin and Marquer (2009). Unfortunately, it is impossible to differentiate the causes of individual differences in the width of the CIs with the current data. We recommend that individual differences in trial-to-trial variability near asymptotic levels of performance are given greater attention in future research (cf. Shing, Schmiedek, Lövdén, & Lindenberger, 2012).

The lack of identifiability of memory search processes with data from a single session may explain why it has not been possible so far to associate different RT patterns with individual differences in reported strategy use (Corbin & Marquer, 2009; see also Marquer & Pereira, 1990) or differential patterns of neural activity (Pelosi et al., 1995). Thus, an important question for future research will be whether observed individual differences in RT patterns, strategy use, and neural activation can be integrated meaningfully on the individual level if sufficient reliability is ensured for the identification of the relevant information (cf. Miller, 2009; Miller et al., 2012; Miller et al., 2009; Miller et al., 2002).

Taken together, our results imply that the cognitive processes that are active during memory search can change within individuals over time and differ between individuals at any given point in time. As a consequence, single-session data may be an insufficient empirical basis for inferring cognitive processes that can be safely assumed to apply to all individuals (of an intended population), and do not necessarily provide a viable route for the identification of cognitive processes at the individual level.

### Are ‘Exhaustive’ and ‘Self-terminating’ Memory Searchers Different?

As the majority of participants could be subsumed under two groups, we examined whether individuals complying with either the exhaustive (*n* = 13) or self-terminating (*n* = 13) memory search model at asymptotic performance differed in assessed covariate measures or parameters of the Sternberg paradigm (Table 7). Notably, consistent with Sternberg’s notion (1975) that self-terminating search is a less efficient memory search process (see also Clifton & Birenbaum, 1970; Corballis & Miller, 1973; Klatzky & Atkinson, 1970), reliable group differences were found in the Reverse Span task (*d* = 1.00; average memory span difference of one item) and in the average *no* response slopes at asymptotic performance (*d* = −0.95; 51 vs. 32 ms/item). Furthermore, group differences in memory span were consistent across all assessed tasks (Digit Span, WM Sorting, Operation and Symmetry Span). On the other hand, no reliable group difference was found in the Digit Symbol task (*d* = −0.15), which has been associated with incidental learning (cf. Joy, Kaplan, & Fein, 2004), and supports the notion that self-terminating search is related to less efficient search and retrieval processes rather than substantial differences in the encoding of information.

However, in contrast to any strong conclusions about memory search efficiency derived from the group statistics, memory spans and memory scanning rates were largely overlapping across groups (Figure 10). An interesting notion that may account for this observation is the concept of vicariance (Lautrey, 2003). It assumes that in principle a set of alternative, substitutable processes may exist that can serve the same general function in the execution of a task; specific processes may arise on the basis of individual differences at the various levels of genetic predisposition, environmental factors, and their complex interactions during individual development (cf. Edelman, 1987). Thus, even within a group of healthy younger adults of mostly university students, a set of alternative, substitutable processes may exist and fulfill the same cognitive function (cf. Corbin & Marquer, 2009). Whereas alternative processes may appear on average different in their efficiency, for any particular individual and process, the functional outcome (i.e., performance) may well be equivalent with the on average more efficient process. Consistent with this notion, no marked group differences in higher order cognition, as for example in general fluid intelligence were observed (Raven Matrices; *d* = −0.11).

Taken together, within a group of healthy younger adults two subgroups were identified using distinct memory search processes. In line with the literature, self-terminating memory search was associated with less efficient memory search on the group level. On the other hand, the distributions of performance measures largely overlapped across groups. The concept of vicariance well accounts for the existence of substitutable but functionally more or less equivalent processes, and strongly underlines the need for careful consideration of the individual level also in the investigation of ‘basic’ cognitive functioning, such as search in memory for digits.

### Generality of Cognitive Processes – Comment on Aggregate Statistics and Replication of Findings

From a methodological perspective, we would like to add a comment on aggregate statistics and a common misinterpretation. As introduced above, aggregate statistics—like Student’s *t*-test or analysis of variance (ANOVA)—fundamentally ignore the individual; the variance introduced by the individuals is treated as error (cf. Gigerenzer, 1987a; Nesselroade, 2010). In this sense, individual differences are rarely in the equation of experimental cognitive psychology, consistent with Cronbach’s conclusion that “individual differences have been an annoyance rather than a challenge to the experimenter” (Cronbach, 1957, p. 674).

Ironically, the proper implication for many commonly used statistics, namely that statistically significant results indicate the existence of some effect at the group level, is often taken as good evidence for the generality of an effect in the sense that the effects hold for individual members of the group. This inference is logically invalid, as the aggregate does not contain information about individual, unless strong additional assumptions are made (cf. Bakan, 1966; <u>Caramazza, 1986</u>; <u>McCloskey & Caramazza, 1988</u>). At best, a few individuals that do not follow the mean patterns may simply go unnoticed; at worst, statistically reliable mixtures of processes may be observed, and the average may not adequately capture even a single individual. Within the present study, the aggregate slope ratio at asymptotic performance provides a characteristic example for this issue.

In the present study, we replicated Sternberg’s original finding in session one. We then noted a substantial change in the mean model as a function of practice, along with substantial individual differences throughout the entire experiment. This pattern of results also informs current debates on the relationship between replicability and generalization in psychology (e.g., Asendorpf et al., 2013). Replication of a mean model—as we have found in the first session— does not guarantee generality of an effect, process, or function at the individual level. Accordingly, replicability is only a necessary but not sufficient condition for generality and must not be used as a substitute for generality.

Taken together, the only way to find out about the generality of a cognitive process is to observe and test it at the individual level. Neither statistical significance nor replication at the aggregate level ensures the existence and generality of psychological phenomena. There have been times in psychology in which the careful observation of individual behavior played a prominent role. The early psychophysics of Wundt, Fechner, and Ebbinghaus, and the careful documentation of learning trajectories during the behaviorist era (e.g., Skinner, 1938) are prominent examples. Nowadays, mathematical psychologists have repeatedly warned against the presence of unnoticed heterogeneity in aggregate data (cf. Luce, 1995). Consequently, we should operate more often in a scientific mode according to which each individual serves as a replication, and adds information about the generality of an effect, function, or process (cf. Danziger, 1990; Gigerenzer, 1987a; Nesselroade, 2010; Nesselroade et al., 2007). Finally, the issue of generality is highly relevant when attempting to find the neural correlates of cognitive processes. When individuals differ among each other in the cognitive processes they use to solve a task, or when a given individual solves a task in more than one way from trial to trial, then the aggregate neural response, either across or within persons, will represent a mixture of neural processing configurations. In this sense, the appreciation of heterogeneity and variation is a necessary challenge for both cognitive psychology (cf. Healey & Kahana, 2013; Lautrey, 2003; Luce, 1995; Siegler, 1987) and cognitive neuroscience (cf. Edelman, 1987; Miller, 2009).

### Summary, Conclusions, and Outlook

This study examined whether the cognitive process model of serial exhaustive search from memory in the Sternberg paradigm inferred from group data held, and was meaningful, at the level of the individual. The model of serial exhaustive search provided an excellent rendition of the first session’s aggregate data but reliability within a single session was insufficient to differentiate between alternative memory search models at the individual level. In line with theoretical considerations, reliability increased substantially when pooling data near stable asymptotic performance. Consequently, cognitive processes became more easily identifiable at the individual level, and individual differences observed near the asymptote could no longer be dismissed as noise and error. At asymptote, the model of serial exhaustive search failed to capture the aggregate data and adequately represented only 13 out of 32 participants; serial self-terminating search consolidated as an equally likely processing mode, and both models together provided a viable description of the majority of individuals. We conclude that the group-based analysis of single-session data yielded a picture of process generality that does not correspond to the processing realities of individual people and their performance when being acquainted with the task.

The current report adds an important case to previously described failures of group analyses to inform about cognitive functioning at the individual level (e.g., Anderson & Tweney, 1997; Estes, 1956; Hayes, 1953; Healey & Kahana, 2013; Siegler, 1987). At the same time, it complements previous work by emphasizing the importance of reliability of measurements and demonstrating that the reliability—as a key prerequisite for the identification of cognitive processes at the individual level—can be improved in a principled way when accounting for the methodological and conceptual issues involved in the repeated assessment of a task.

As Wagenmakers and Waldorp (2006) have pointed out, the “main objective of the scientific enterprise is to find explanations for the phenomena we observe” (p. 99). Theories in mathematical psychology (Luce, 1995), developmental psychology (P. B. Baltes, Lindenberger, & Staudinger, 2006; Lautrey, 2003; Li & Lindenberger, 2002), and neuroscience (Edelman, 1987) have convincingly argued that the mapping of processes onto behavior may not be unitary but variable within and between persons. From this perspective, it seems mistaken to search for a one-to-one correspondence between a cognitive process and an experimental paradigm; instead, the more meaningful and realistic goal is to investigate which individuals implement what process under a given set of contextual constraints (cf. Dhami et al., 2004). In this sense, it is essential to put strong efforts into describing as accurately as possible the full extent of phenomena at the level of the individual that await their explanation. An important challenge along this way will be the acquisition of data that allows unequivocal identification of psychological processes at the individual level.

## Appendix A

### Covariate Session

The *personal and health questionnaire* included questions regarding educational level, current occupation, health status (with an extensive set of questions ensuring suitability of participants for EEG recordings and MRI acquisition), handedness scale (after Oldfield, 1971), and CES-D depression scale (Radloff, 1977). The *daily questionnaire* will be described below.

*Visual acuity* was assessed by the computerized “Freiburg Visual Acuity test” (Bach, 1996, 2007). *Digit Span* and *Reverse Span* were adopted from the Wechsler Adult Intelligence Scale (WAIS; Wechsler, 1958). *Number sorting* is similar to the Digit and Reverse Span task; between three and eight numbers from 1 to 12 where read aloud to the participant with approximately one number per second; participants had to report subsequently the set of numbers in ascending order. *Digit Symbol Substitution* task was again adopted from the WAIS. *Raven Matrices* were 18 selected items from the *Raven’s Advanced Progressive Matrices*. *Operation Span* tasks were adopted from the automated version provided by Unsworth and colleagues (Unsworth, Heitz, Schrock, & Engle, 2005). In short, participants are presented alternately with simple mathematical equations in the form of “(8 / 2) − 1 = ?” and consonants; after two to seven presentations they are required to recall the consonants in the order of presentation (serial recall). We generated an alternative version, where participants were alternately presented with three consonants and had to indicate whether they were in alphabetical order, and with single digits; after two to seven alternations they had to recall the digits in the order of presentation. Finally, the *Symmetry Span* task (Unsworth et al., 2005) was conducted as described by Kane and colleagues (Kane et al., 2004).

### Repeated (EEG and Practice) Sessions – Questionnaires and Tasks

Questionnaires and cognitive tasks were identical for EEG and practice sessions. A *daily questionnaire* in the beginning of each session was adapted from the questionnaire used in the COGITO study (Schmiedek, Lövdén, & Lindenberger, 2010). In short, it assessed negative affect, daily stressors, unspecific and stressor-related intrusive thoughts (see Brose, Schmiedek, Lövdén, & Lindenberger, 2011, for a detailed description), sleep in the night preceding the experimental session, general health status, consumption of alcohol, caffeine, and drugs (pharmaceuticals) within the last 24 h preceding the experimental session, and overall motivation to complete the cognitive tasks. The *second daily questionnaire* was provided at the end of each session, asking for disturbances during the session, attitude towards and motivation regarding the cognitive tasks during the just completed experimental session.

Besides the Sternberg task and the strategy questionnaire, in every session choice reaction, number comparison, and visually presented digit and reverse span tasks were employed. In the *Choice Reaction* task, the letters *R* and *L* were presented equivalent to the Sternberg task (i.e., at the same location on the screen with the same font type and size) in random order; participants had to indicate as fast and accurately as possible which letter had appeared on the screen by pressing a button with their right index finger for *R* and with their left index finger for *L*, respectively. In the *Number Comparison* task, participants were presented with two strings of three to six digits besides each other; participants had to indicate as fast and accurately as possible whether the strings were the same or not by pressing a button with their right index finger for equivalent and with their left index finger for different strings of digits; strings deviated in maximally one of the digits. The *Digit* and *Reverse Span* task in the repeated sessions deviated from the WAIS version used in the covariate session; digits were presented visually, exactly in the same way as in the Sternberg task. After presentation of the last digit, a warning tone indicated the beginning of the recall; participants were then required to recall the digits in the order of presentation, and the correctness of the serial recall was recorded. Eight different sets of random digit strings were generated for the eight repeated sessions.

### EEG Recordings

In session one, seven, and eight EEG was concurrently recorded with the Sternberg and the choice reaction task. Preparation of the EEG electrodes took approximately one hour. In addition to the behavioral sessions, EEG resting state recordings were acquired before the tasks were conducted. 2 min with eyes closed and 2 min with eyes open were recorded, while subjects were asked to sit as still and relaxed as possible and fixate a fixation cross during the eyes open condition (cf. Grandy et al., 2013). In addition, recordings were made where participants were asked to alternately open and close their eyes every 5 sec in response to a beep tone (cf. Whitten, Hughes, Dickson, & Caplan, 2011). Finally, visual evoked potentials (cf. Odom et al., 2004) with checkerboards of two different pattern sizes were recorded.

## Appendix B

### Linearity of Slopes

Within MLM, we tested formally for quadratic trends in RTs as a function of set size by introducing quadratic fixed effects for negative and positive slopes, respectively. For the first session, likelihood ratio test indicated no significant increase in model fit when adding quadratic fixed effects [Δ*χ*^2^(2) = 1.98, *p* = .371; quadratic trend in positive slopes only: Δ*χ*^2^(1) = 1.85, *p* = .173; quadratic trend in negative slopes only: Δ*χ*^2^(1) = 0.13, *p* = .723]. Thus the linear model sufficiently accounted for the overall linear increase in RTs as a function of set size. Data for sessions two to eight are reported in Table 8.

**Table 8.**
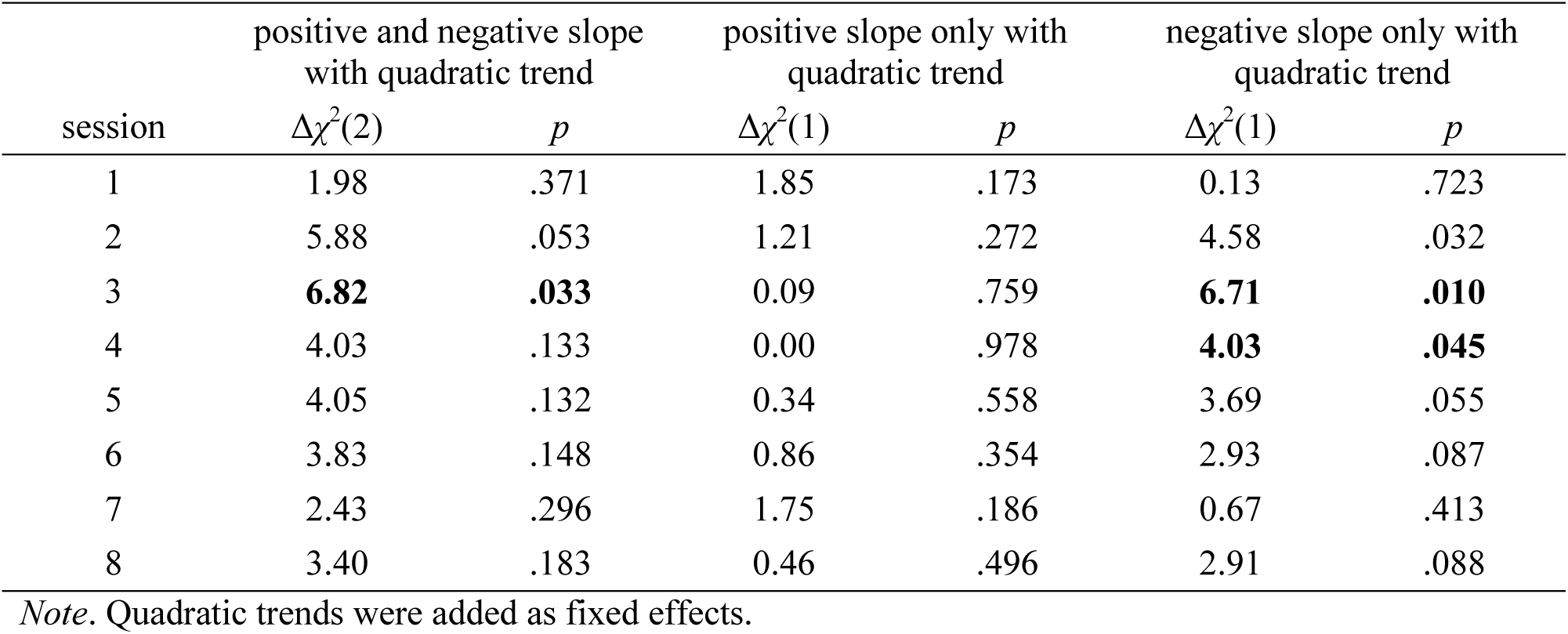
(Appendix B). Testing the reliability of mean quadratic trends in addition to linear trends within MLM.

| session | positive and negative slope<br>with quadratic trend |  | positive slope only with<br>quadratic trend |  | negative slope only with<br>quadratic trend |  |
| --- | --- | --- | --- | --- | --- | --- |
| | $\Delta\chi^2(2)$ | $p$ | $\Delta\chi^2(1)$ | $p$ | $\Delta\chi^2(1)$ | $p$ |
| 1 | 1.98 | .371 | 1.85 | .173 | 0.13 | .723 |
| 2 | 5.88 | .053 | 1.21 | .272 | 4.58 | .032 |
| 3 | **6.82** | **.033** | 0.09 | .759 | **6.71** | **.010** |
| 4 | 4.03 | .133 | 0.00 | .978 | **4.03** | **.045** |
| 5 | 4.05 | .132 | 0.34 | .558 | 3.69 | .055 |
| 6 | 3.83 | .148 | 0.86 | .354 | 2.93 | .087 |
| 7 | 2.43 | .296 | 1.75 | .186 | 0.67 | .413 |
| 8 | 3.40 | .183 | 0.46 | .496 | 2.91 | .088 |

*Note.* Quadratic trends were added as fixed effects.

## Footnotes

1 Comparison specifically of *no* responses has been conducted under the assumption that in order to initialize a *no* response memory has to be searched ‘exhaustively’ within both groups (cf. Sternberg, 1975).

## Notes

Author Note: This study was conducted within the project “*Cognitive and Neural Dynamics of Memory Across the Lifespan (CONMEM)”* at the Max Planck Institute for Human Development, Berlin, Germany. The work was supported by the Max Planck Society and the Gottfried Wilhelm Leibniz Prize 2011 of the German Research Foundation awarded to UL. MW-B’s work is supported by a grant from the German Research Foundation (DFG, WE 4269/3-1) as well as an *Early Career Research Fellowship 2017 – 2019* awarded by the Jacobs Foundation. We thank Lene-Marie Gassner and all research assistants involved in data collection.

### Competing Interest Statement

The authors have declared no competing interest.

### Summary of Updates

As discussed in an email-conversation (Re: BIORXIV/2017/126490 - Werkle-Bergner; February, 12, 2025), the revision just removed one sentence from the title page. Otherwise the manuscript stayed unchanged.

